# Vagus nerve stimulation increases stomach-brain coupling via a vagal afferent pathway

**DOI:** 10.1101/2021.10.07.463517

**Authors:** Sophie Müller, Vanessa Teckentrup, Ignacio Rebollo, Manfred Hallschmid, Nils B. Kroemer

**Author notes:** **Corresponding author** Dr. Nils B. Kroemer, Calwerstr. 14, 72076 Tübingen, Germany, Twitter: @cornu_copiae.

## Abstract

Maintaining energy homeostasis is vital and supported by vagal signaling between digestive organs and the brain. Previous research has established a gastric network in the brain that is phase synchronized with the rhythm of the stomach, but tools to perturb its function were lacking. Here, we investigated the effect of acute right-sided transcutaneous auricular vagus nerve stimulation (taVNS) versus sham stimulation (randomized crossover-design) on stomach-brain coupling. In line with preclinical research, taVNS increased stomach-brain coupling in the nucleus of the solitary tract (NTS) and the midbrain while boosting coupling across the brain. Crucially, in the cortex, taVNS-induced changes in coupling occurred primarily in transmodal regions and were associated with changes in hunger ratings as indicators of the subjective metabolic state. Hence, taVNS alters stomach-brain coupling via an NTS-midbrain pathway that signals gut-induced reward, potentially paving the way for novel treatments in disorders such as Parkinson’s disease or depression.

**Graphical Abstract:** 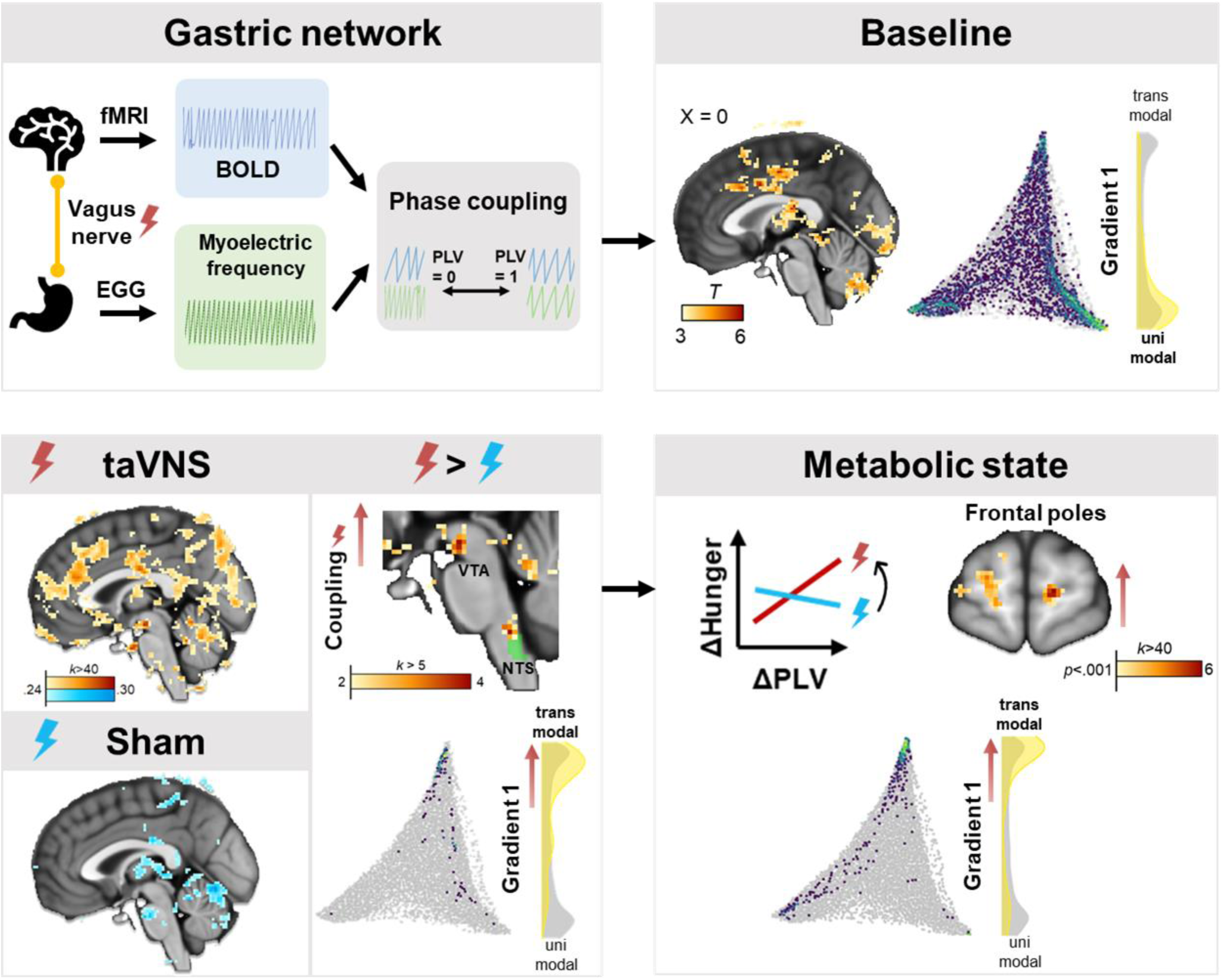

**Highlights:**

- Transcutaneous vagus nerve stimulation (taVNS) can emulate interoceptive signals
- taVNS boosts stomach-brain coupling in the brainstem, midbrain, and transmodal cortex
- taVNS-induced changes in stomach-brain coupling mirror subjective hunger ratings

## Introduction

Communication between the gut and the brain is central for maintaining energy homeostasis (Waterson and Horvath, 2015). Interoceptive signals from peripheral organs involved in energy metabolism, such as the stomach, convey the physiological state of the body via vagal afferents to the nucleus of the solitary tract (NTS) in the brainstem (Holtmann and Talley, 2014; Powley et al., 2011), thereby constantly updating homeostatic information in the brain. Afferent vagal signaling from the stomach originates from chemo- and mechanoreceptors located in the stomach wall and has been shown to mediate hunger and satiety (Folgueira et al., 2014). Efferent vagal signaling modulates digestion by regulating the movement of the gastrointestinal smooth muscles directly or via the interstitial cells of Cajal (Lundgren, 1983), the gut pacemaker cells. However, it is still largely unknown how the brain processes information conveyed by the stomach and whether afferent vagal stimulation can alter the communication between the stomach and the brain in humans.

Previous research on the stomach-brain axis has established the existence of a gastric network in the brain, whose activity is coupled to the 0.05 Hz myoelectrical rhythm intrinsically produced in the stomach that paces digestive contractions (Rebollo et al., 2018). Interoceptive signaling has been hypothesized to entrain perceptual and attentional processes on the neural level (Rassi et al., 2019); this has been demonstrated for cardiac (Allen et al., 2019) and respiratory (Varga and Heck, 2017) rhythms. Rhythmic signals originating from the stomach and mediated by the vagus nerve have been linked not only to appetite (Hussain and Pan, 2009; Mattes et al., 2019), but also motivation (Alhadeff and Grill, 2014; Kanoski et al., 2013; Nord et al., 2021), memory formation (Mandal et al., 2018; Suarez et al., 2018) and affect (Mayer, 2011). Consequently, a widespread cortical network relying on a variety of neuromodulatory systems is coupled to the gastric rhythm (Rebollo et al., 2021), suggesting a role for gastric network oscillations in relevant neuromodulatory and behavioral functions. Furthermore, dysfunctions of this network might be associated with disorders such as Parkinson’s disease (Heimrich et al., 2019; Svensson et al., 2015) that is characterized by aberrant vagal signaling from the gut and motivational impairments (Mazzoni et al., 2007), both of which might be mechanistically linked (Breen et al., 2019).

However, to better characterize the functional role of the gastric network, causal manipulations of stomach-brain axis signaling are necessary, a task that has so far remained elusive. Transcutaneous auricular vagus nerve stimulation (taVNS) has been shown to afferently increase activity in the NTS (Frangos et al., 2015; Sclocco et al., 2019; Teckentrup et al., 2021; Yakunina et al., 2017) and improve motivation (Neuser et al., 2020), memory (Giraudier et al., 2020; Jacobs et al., 2015; Vazquez-Oliver et al., 2020), and mood (Ferstl et al., 2021; Kraus et al., 2007). Vice versa, gastric motility was modulated by manipulations of an efferent vago-vagal pathway (Hong et al., 2019; Steidel et al., 2021; Teckentrup et al., 2020). Here, we used taVNS with concurrent electrogastrography (EGG) and fMRI to assess the effect of vagal afferent activation on stomach-brain coupling in healthy individuals (Fig. 1A). We expected taVNS to increase gastric coupling in regions with known vagal afferent projections, such as the NTS. After corroborating the existence of the gastric network at baseline, we found that taVNS increases stomach-brain coupling in the NTS and the dopaminergic midbrain as well as in transmodal cortical regions, where changes in coupling are correlated with changes in interoceptive ratings of hunger (vs. satiety). These results show that stomach-brain coupling can be modulated with taVNS via a pathway that conveys gut-induced reward signals.

**Figure 1:**
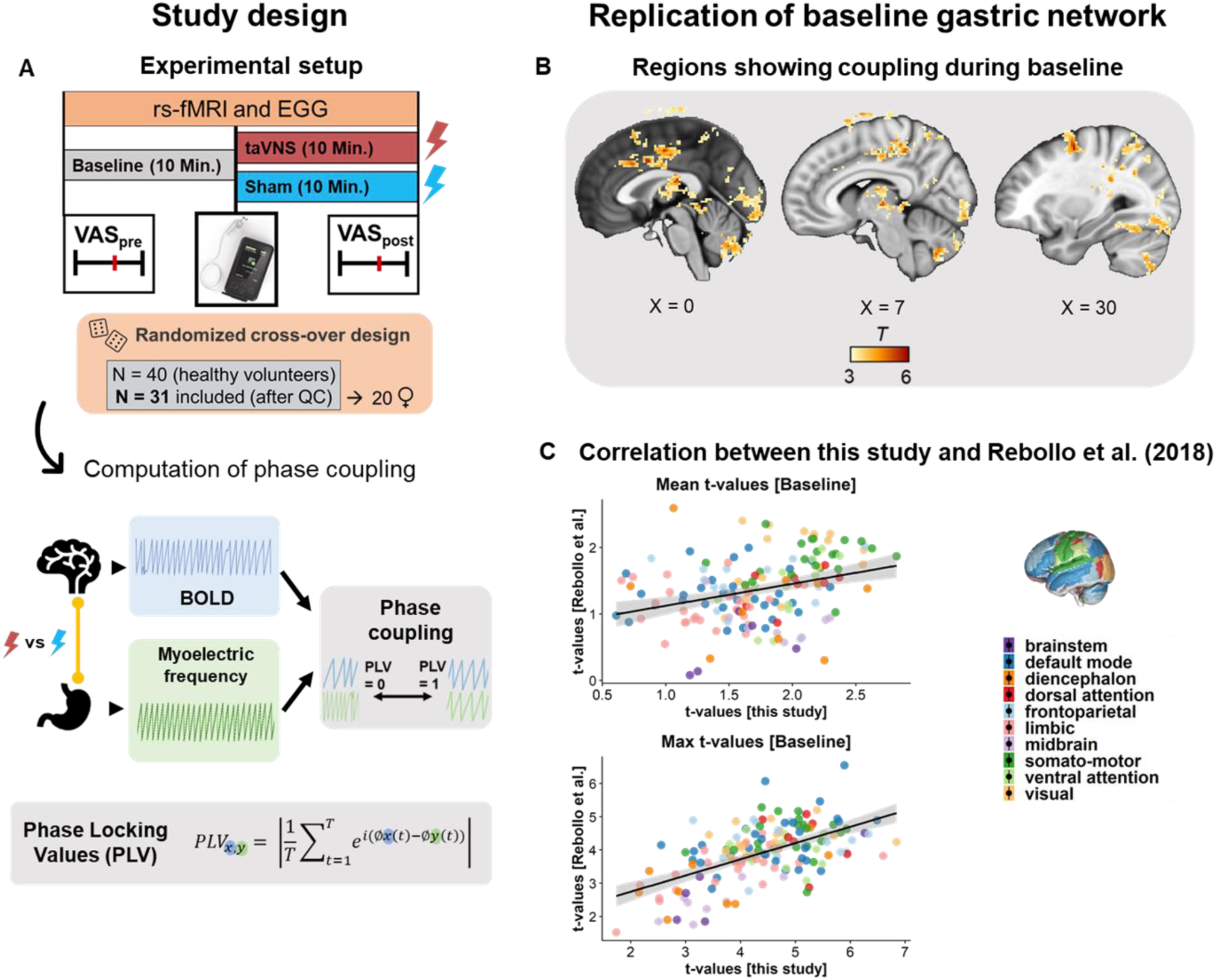
Study design and corroboration of the gastric network during baseline scans (before stimulation). **A**: We measured resting-state fMRI and electrogastrography (EGG) in 31 healthy participants (20 women) for 10 min of baseline and 10 min during stimulation (taVNS vs. sham, collected on different days). The coupling between the stomach and the brain was expressed by the phase-locking value (PLV), indicating the synchrony between time series. **B**: We replicated the core regions of the canonical gastric network at baseline by observing coupling in the postcentral gyrus, the cingulate gyrus, the precuneus, the occipital cortex, the fusiform gyrus, the inferior frontal gyrus, the inferior and superior parietal lobe, the thalamus, and the inferior cerebellum. **C**: Within anatomically defined brain regions, we observed a moderate to high correlation between PLVs obtained in our sample and the independent sample of Rebollo et al. (2018), average *t*-values: *r* = .318, *p* < .001, maximal *t*-values: *r* = .57, *p* < .001, *t*-values available in Table S1.

## Results

We analyzed phase coupling between the fMRI BOLD signal (providing high spatial resolution in deep brain regions) and the myoelectric signal of the stomach using the phase-locking value (PLV), a measure reflecting the synchrony between two time series. We successfully replicated the anatomical and functional aspects of the gastric network as described by Rebollo *et al*. (2018) during baseline (Fig. 1, t-values in Table S1, all maps available here: https://neurovault.org/collections/QNGZBQGF/) indicating that stomach-brain coupling is a robust phenomenon.

### taVNS boosts stomach-brain coupling in the NTS, midbrain, and transmodal cortex

To evaluate whether stomach-brain coupling can be enhanced by taVNS, we compared PLV maps during taVNS and sham stimulation (relative to baseline) using a full-factorial model (i.e., Stimulation × Time interaction). Within our a priori regions of interest, we found that taVNS increased stomach-brain coupling in the NTS (Fig. 2A; *p*_svc_ = .002, *t*_*max*_ = 4.33) which is the first entry point of vagal afferents in the brain. Moreover, we observed increases in the dopaminergic midbrain (Fig. 2A; combined mask VTA & SN: *p*_svc_ = .037, peak in VTA: *t*_*max*_ = 3.40, *p*_svc_ = .006). At a whole-brain level (corrected for multiple comparisons via cluster extent), we observed that taVNS increased stomach-brain coupling in transmodal cortical regions, namely in the mid frontal gyrus (Fig. 2B; *t*_*max*_ = 4.97, *k* = 63) and the precuneus (*t*_*max*_ = 4.25, *k* = 54). Also, we found a significantly increased interindividual variance in stomach-brain coupling during taVNS versus sham in several regions (*p* < .01, *F*(30,30) > 2.39), such as the anterior cingulate cortex (Fig. 2B; *F*(30,30) = 2.96, *p* = .002), the left secondary somatosensory cortex (*F*(30,30) = 2.94, *p* = .002), the right insular cortex (*F*(30,30) = 2.66, *p* = .005), and the right supramarginal gyrus (*F*(30,30) = 2.52, *p* = .007), indicating successful taVNS-induced perturbation.

**Figure 2:**
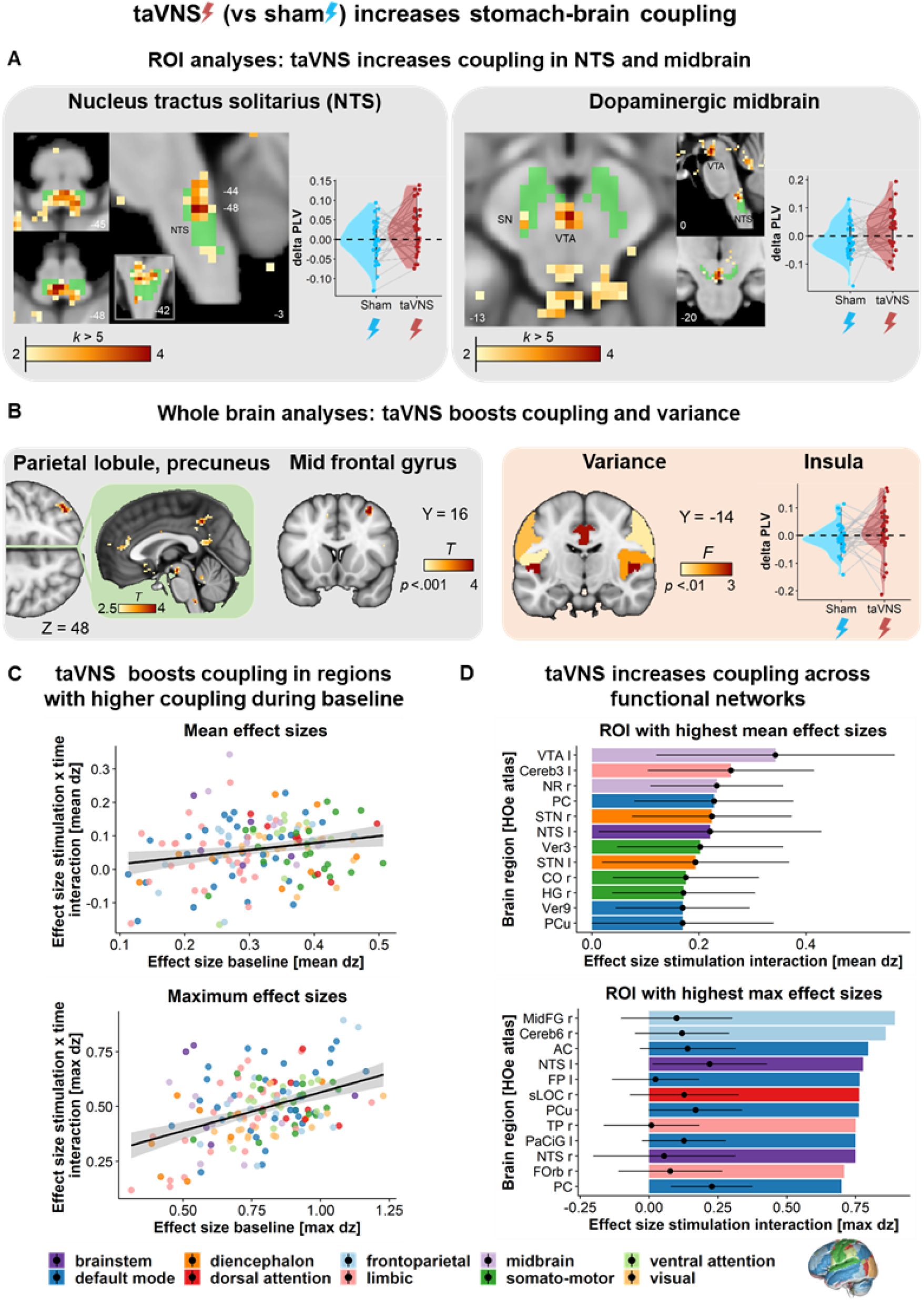
Transcutaneous vagus nerve stimulation (taVNS) increases stomach-brain coupling. **A**: taVNS boosts coupling in key target regions of vagal afferents: the NTS (*t*_*max*_ = 4.33, *p*_*svc*_ = .002) and the dopaminergic midbrain (peak in the VTA/SN: *p*_svc_ = .037, *t*_*max*_ = 3.40). **B**: taVNS increases coupling in the precuneus/superior parietal lobe (*t*_*max*_ = 4.25, *p*_*FWE-corr cluster-level*_ = .041) and the mid frontal gyrus (*t*_*max*_ = 4.97, *p*_*FWE-corr cluster-level*_ = .019), as well as variance in PLV changes in several regions (vs. sham), such as in both insular cortices and the anterior cingulate (AC) cortex. See also Figure S1. **C**: taVNS boosts coupling in regions that are intrinsincally coupled with the stomach at baseline as indicated by Pearson correlations between baseline and stimulation interaction effects: (mean *d*_*z*_ over each region: *r* = .21, *p* = .009; maximum *d*_*z*_ within each region: *r* = .44, *p* < .001). **D**: Selected regions with the highest effect sizes, calculated voxel-wise (error bars show M ± SD). VTA: ventral tegmental area, Cereb: cerebellum, NR: nucleus. ruber, PC: posterior cingulate, STN: subthalamic ncl., NTS: nucleus tractus solitarius, Ver: vermis, CO: central opercular cortex, HG: Heschl’s gyrus, PCu: precuneus, MidFG: mid frontal gyrus, sLOC: superior lateral occipital cortex, TP: temporal pole, FOrb: frontal orbital cortex.

To assess potential influences of taVNS strength on stomach-brain coupling, we calculated the correlation between taVNS strength and changes in PLV (taVNS - sham), again using a factorial model in SPM. We found strong positive correlations in both insular cortices (Fig. S1, maps on NeuroVault; right: *t*_*max*_ = 4.52, *k* = 102; left: *t*_*max*_ = 4.10, *k* = 190) and in the right supramarginal gyrus (*t*_*max*_ = 4.61, *k* = 74). Crucially, all three regions showed a significantly increased variance during taVNS, indicating that interindividual differences in stimulation strength (matched according to sensory aspects) might contribute to the observed variance. Still, gastric myoelectric frequency did not change during taVNS in our sample (Teckentrup *et al*., 2021), indicating that changes in stomach-brain coupling might precede changes in gastric motility that occur after prolonged stimulation (Teckentrup *et al*., 2020).

To evaluate whether taVNS increases coupling in regions that are intrinsically coupled at baseline, we computed the correlation between the effect sizes (*d*_*z*_) of each region for the baseline (before taVNS and sham) and for taVNS-induced changes (taVNS – sham) in PLVs. Both mean effect sizes (*r* = .21, *p* = .009), and maximum effect sizes within each region (*r* = .44, *p* < .001) were significantly correlated, demonstrating that taVNS boosted stomach-brain coupling in regions with higher intrinsic coupling at baseline (Fig. 2C, Table S2). However, the largest taVNS effects on PLVs were observed in regions outside the baseline gastric network, notably in the frontoparietal network, the default-mode network (DMN), the midbrain, and the brainstem (Fig. 2D, networks assigned according to Yeo et al. (2011)). Analogous to Rebollo *et al*. (2021), we also analyzed the hierarchical distribution of the taVNS effect on the gastric network along the first two cortical resting-state gradients as described by Margulies et al. (2016). These gradients characterize resting-state networks in terms of a cortical functional hierarchy, where the first gradient extends from uni- to transmodal (with two peaks at the extremes) and the second gradient from visual to somatomotor-auditory regions (Fig. 3A). In line with Rebollo et al. (2021), the baseline gastric network was primarily associated with unimodal cortical regions (Fig. 3B; *p* < .001 deviation from the standard resting-state gradient, Kolmogorov-Smirnov test). Intriguingly, taVNS shifted this distribution towards transmodal cortical regions, such as the DMN and the frontoparietal network (Fig. 3C; *p* < .001).

**Figure 3:**
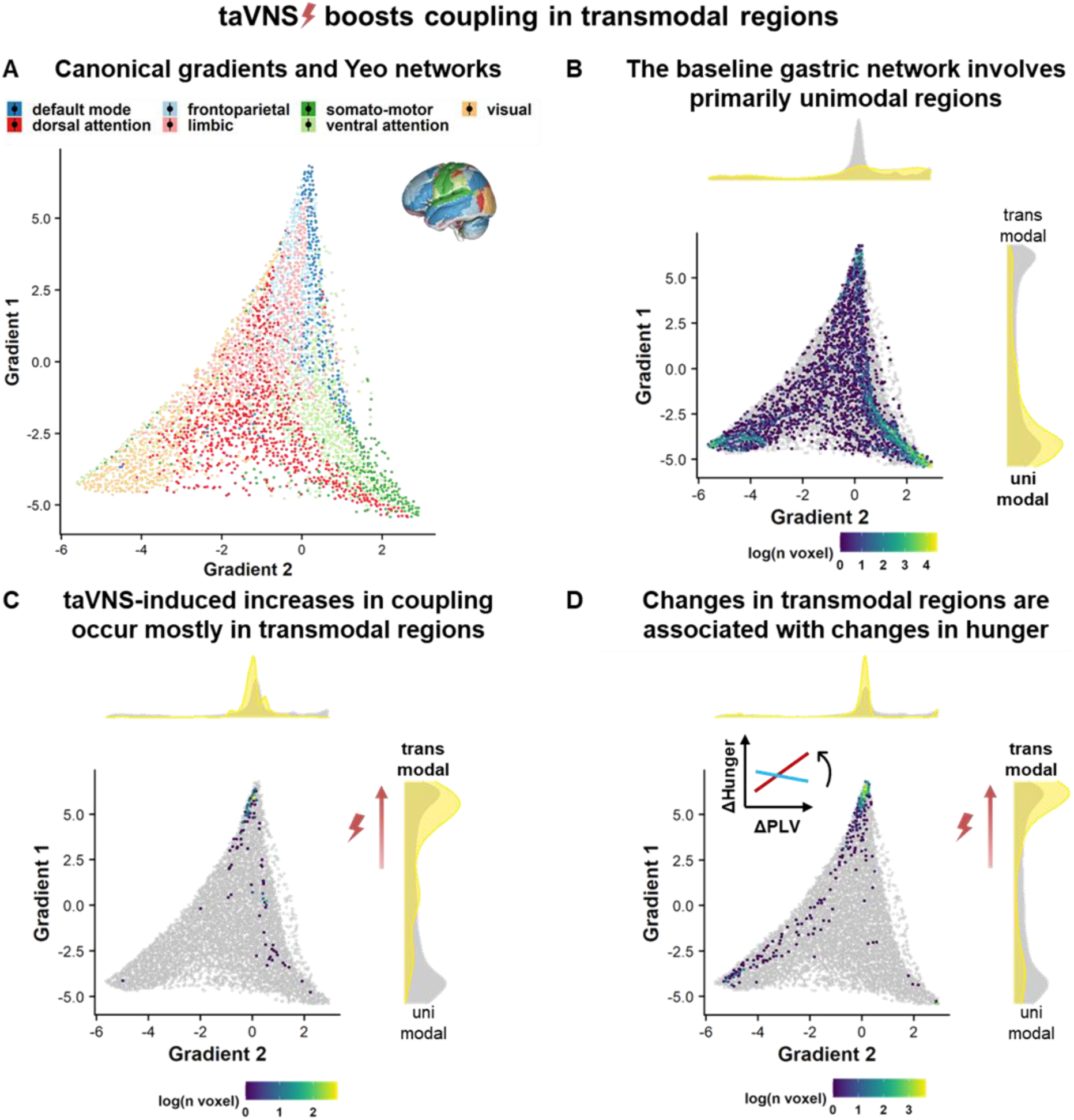
Non-invasive vagus nerve stimulation (taVNS) primarily boosts coupling in transmodal cortical regions. **A**: Distribution of the cortical gradients described by Margulies *et al*. (2016). Each dot depicts a bin of voxels (100 bins per gradient) and was color-coded according to the predominant Yeo network in each bin. Gradient 1 extends from uni- to transmodal regions while gradient 2 extends from visual to somato-motor-auditory regions. **B**: The plot compares the distribution of voxels that show significant taVNS-induced changes in coupling (highlighted) to the standard cortical gradients described by Margulies *et al*. (2016) depicted in grey. The colors indicate the numbers of voxels per bin. In line with Rebollo et al. (2021), the voxels of the baseline gastric network belonged primarily to unimodal regions, such as visual and somato-motor areas. **C**: In contrast, taVNS increased stomach-brain coupling primarily in transmodal regions, such as the default mode network and the frontoparietal network, which is in line with an increase in parasympathetic modulation. **D**: This plot shows the distribution of voxels that show a significant correlation between taVNS-induced changes and sensed changes in hunger (highlighted) compared to the standard cortical gradients described by Margulies *et al*. (2016) depicted in grey. Intriguingly, taVNS-induced changes in stomach-brain coupling were correlated with subjectively sensed changes in hunger mostly in transmodal cortical regions as well, suggesting that changes in stomach-brain coupling in transmodal cortical regions may play a role in sensing interoceptive signals related to hunger (i.e., metabolic state).

### taVNS strengthens the link between changes in stomach-brain coupling and changes in hunger

After demonstrating that taVNS boosts stomach-brain coupling, we analyzed whether changes in subjective ratings of metabolic state (assessed before and after the measurement) were associated with individual differences in taVNS-induced changes. To better evaluate these changes, participants were instructed to come to the lab neither hungry nor full and we defined subjective metabolic state as the difference between visual analog scale (VAS)-rated hunger (“How hungry are you?”) and satiety (“How sated are you?”), with positive values corresponding to increased hunger and negative values corresponding to increased satiety. These state ratings showed an increase in hunger during the session (*p* < .001) that emerged with considerable individual variability but did not differ depending on the stimulation condition (Fig. 4A; *p* = .482, mixed-effects model). However, we found that taVNS increased the correlation of changes in stomach-brain coupling with changes in hunger ratings in regions across the cortex (Fig. 4B-E, whole-brain corrected), such as the frontal poles (right: *t*_*max*_ = 6.49, *k* = 40; left: *t*_*max*_ = 5.16, *k* = 137), the cerebellum (*t*_*max*_ = 5.14, *k* = 200), the fusiform gyrus (*t*_*max*_ = 4.63, *k* = 92), the precuneus (left: *t*_*max*_ = 4.54, *k* = 100; right: *t*_*max*_ = 4.49, *k* = 67), the inferior parietal lobe (*t*_*max*_ = 4.25, *k* = 60), and the declive (*t*_*max*_ = 4.41, *k* = 55, Table S3, maps on NeuroVault). In line with the main effect of taVNS, the correlation of taVNS-induced changes in PLV with subjective hunger ratings showed a shift towards transmodal cortical regions (Fig. 3D; *p* < .001 compared to the cortical gradients by Margulies *et al*. (2016)). Collectively, these findings support the idea that taVNS-induced changes in stomach-brain coupling can be sensed, indicating that emulated vagal afferents may act as interoceptive signals.

**Figure 4:**
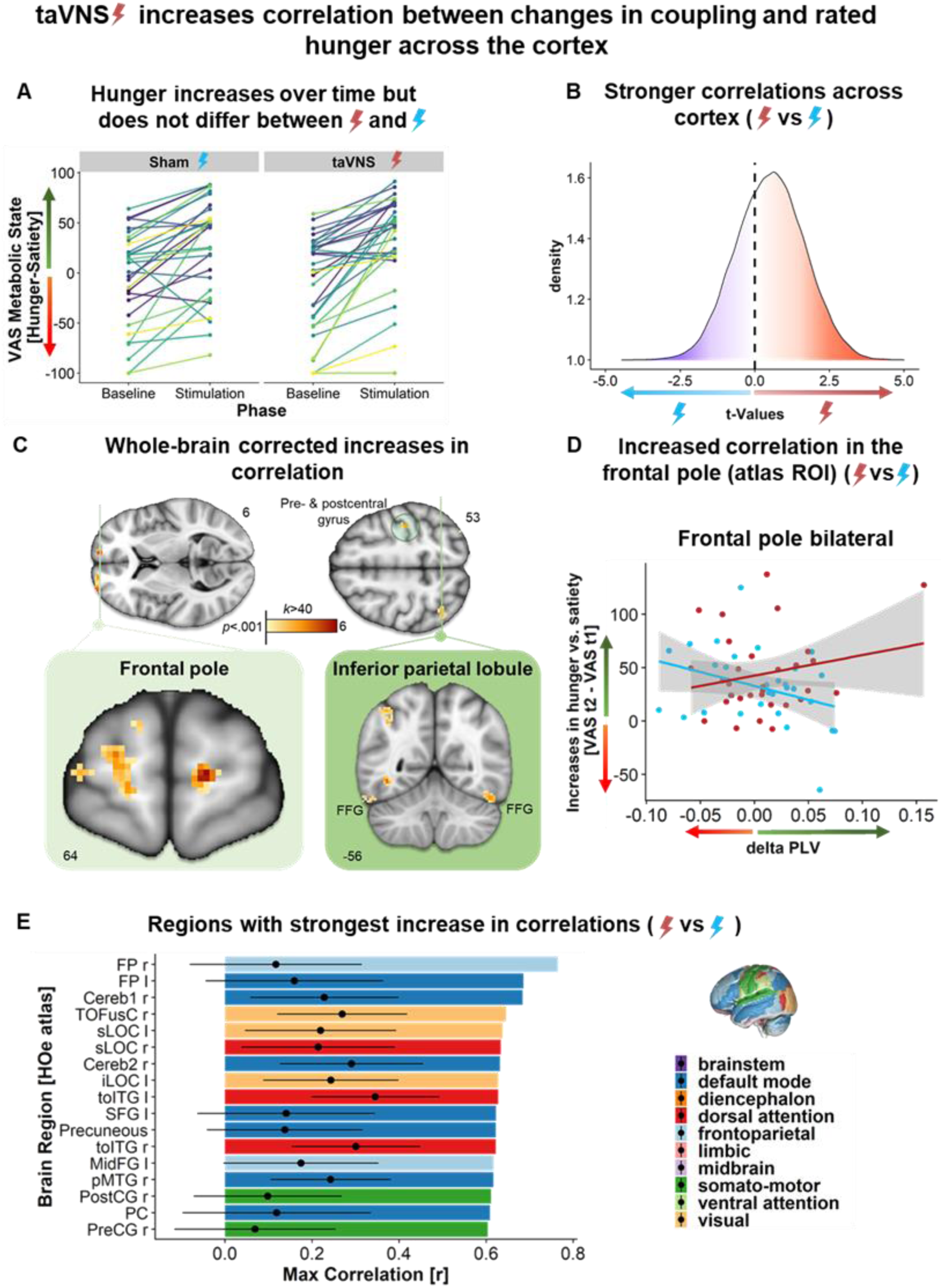
taVNS increases the correspondence of changes in stomach-brain coupling with subjective changes in metabolic state. **A**: Participants became hungrier during the scan (*p* < .001), but there was no interaction with stimulation (*p* = .482). **B**: taVNS increased the correlation between increases in hunger and changes in stomach-brain coupling across the cortex (density plot of the change in correlation (taVNS vs sham) of all cortical voxels, positive values indicate higher correlation). **C**: Clusters of voxels showing significant increases in correlation with subjective changes in metabolic state after whole-brain correction for multiple comparisons: frontal poles (*t*_*max*_ = 6.49), fusiform gyrus (*t*_*max*_ = 4.63), precuneus (*t*_*max*_ = 4.54), and inferior parietal lobe (*t*_*max*_ = 4.25) (Table S3, maps on NeuroVault). **D**: taVNS increases the correlation between PLV changes and metabolic state ratings in the frontal poles (regions from Harvard-Oxford brain atlas). **E**: During taVNS, several regions showed an increased correlation between changes in coupling and metabolic state and the strongest effects were found in the DMN, the visual network, the frontoparietal network, and the dorsal attention network. Correlation was calculated voxelwise over ROI, bar shows maximum r, error bar indicates mean ± standard deviation. [FP: frontal pole, Cereb: cerebellum, TOFusC: temporal occipital fusiform gyrus, sLOC: superior lateral occipital cortex, iLOC: inferior lateral occipital cortex, toITG: temporooccipital inferior temporal gyrus, MidFG: mid frontal gyrus, pMTG: posterior middle temporal gyrus, PostCG: postcentral gyrus, PC: posterior cingulate, PreCG: precentral gyrus]

## Discussion

The recently discovered gastric network in the brain is thought to play a vital role in energy homeostasis (Choe et al., 2021; Rebollo *et al*., 2018; Rebollo *et al*., 2021), but the lack of causal manipulations of the network has so far hampered progress in delineating its functional role in humans. Here, we used non-invasive taVNS to emulate the activation of vagal afferents that are known to regulate the intrinsic rhythm of the stomach via a vago-vagal pathway (Shapiro and Miselis, 1985). For the first time in humans, we show that taVNS applied at the right ear boosts stomach-brain coupling in the NTS and the dopaminergic midbrain, a pattern that strongly resembles the pathway for gut-induced reward via the right nodose ganglion previously described in rodents (Han et al., 2018). Furthermore, we demonstrate increased coupling in transmodal cortical regions, as well as in regions that display high instrinsic coupling under baseline conditions. Crucially, we also show that taVNS increases the correlation between changes in stomach-brain coupling and in subjectively sensed hunger, again primarily in transmodal cortical regions. Consequently, we conclude that taVNS modulates stomach-brain coupling via an NTS-midbrain pathway, while changes in cortical coupling might reflect stronger integration of interoceptive signals encoding the current metabolic state of the body. Collectively, these results point to a vital role of the gastric network in supporting the maintenance of energy homeostasis. Thus, taVNS modulation of stomach-brain coupling might be a promising approach in the treatment of neurological or mental disorders with a high prevalence of gastrointestinal symptoms, such as Parkinson’s disease or depression (Kaut et al., 2019).

### Anatomical pathways for the taVNS effect on the gastric network

Since the NTS is the main entry point of vagal afferents to the brain and is robustly activated by taVNS (Frangos *et al*., 2015; He et al., 2013; Teckentrup *et al*., 2021; Yakunina *et al*., 2017), we expected taVNS to increase stomach-brain coupling in this region as well; this hypothesis was confirmed. The NTS is a crucial part of the gastric-vagal reflex: vagal afferents delivering interoceptive information (e.g., mechanical, hormonal, chemical) from the stomach reach the NTS, get integrated and projected to the dorsal vagal motor nucleus, which then sends efferent commands back to the stomach (Berthoud and Neuhuber, 2000; Neuhuber and Berthoud, 2021; Shapiro and Miselis, 1985; Waise et al., 2018). Previous findings also suggest the involvement of this pathway in the efferent modulation of gastric frequency following vagal stimulation in humans (Hong *et al*., 2019; Steidel *et al*., 2021; Teckentrup *et al*., 2020). Although the anatomical pathways mediating the integration of gastric signals in the brain are not fully resolved (Azzalini et al., 2019), it is known that the NTS projects to the parabrachial nuclei (PBN), where vagal and spinal afferents are integrated. They are then projected further to the thalamus which acts as a relay station to cortical regions of the gastric network (Rebollo and Tallon-Baudry, 2021; Rebollo *et al*., 2021). In addition, the NTS also projects to the dopaminergic midbrain which is involved in neural reward circuits, nutrient sensing, and food-seeking behavior (Fernandes et al., 2020; Han *et al*., 2018). In line with previous work, we found that taVNS boosts stomach-brain coupling in the dopaminergic midbrain, which adds to the growing literature suggesting a role for dopamine in the control of digestion (Anselmi et al., 2017). Modulations of the VTA may also occur indirectly via the PBN inasmuch as connections between the PBN and the VTA have been shown to be involved in appetite and taste perception (Boughter et al., 2019) as well as in somatic and visceral nociception (Dunckley et al., 2005). From a clinical perspective, increased stomach-brain coupling in the VTA/SN provides potential links to prevalent gastrointestinal symptoms in several disorders, such as mood and anxiety disorders (Huang et al., 2021; Mussell et al., 2008; Soderquist et al., 2020) or Parkinson’s disease (PD) (Lubomski et al., 2020; Marrinan et al., 2014; Mrabet et al., 2016; Santos-Garcia et al., 2015), in which the dopaminergic midbrain is severely affected (Aurora et al., 2021). Illustratively, Kaut *et al*. (2019) provided promising evidence that taVNS could improve gastrointestinal symptoms in PD, highlighting the potential of a clinical application, with our findings providing a much-needed mechanistic link that requires further investigation. To summarize, our results suggest a strong involvement of a vagal-midbrain pathway that may facilitate the integration of metabolic signaling into larger brain networks to potentially shape goal-directed behavior to ensure that energetic demands are considered.

### The role of the gastric network in maintaining metabolic balance

Given that feeding and digestion are the vital functions of the digestive system, it is likely that the function of the gastric network is to support the maintenance of long-term energy homeostasis (Azzalini *et al*., 2019; Choe *et al*., 2021; Rebollo *et al*., 2021). Our finding that taVNS increases the correlation between changes in stomach-brain coupling and subjectively sensed changes in metabolic state provides intriguing support for the alleged link between the gastric network and interoceptive signaling. This is in line with the work by Levakov et al. (2021), who have recently shown that activity in the gastric network is related to future weight loss and argue for the involvement of the gastric network in the integration of metabolic signals to control body weight. We found that transmodal regions show the highest increase in correlation between changes in coupling and sensed hunger (vs. satiety) during taVNS. This possibly reflects the hierarchically overarching integration of vital metabolic feedback, buffered in unimodal cortical areas, into larger brain networks in response to vagal afferent activation. The strong involvement of visual regions might reflect increased visual attention to (potential) food at times of increased hunger, which would fit the alleged importance of the gastric network for energy homeostasis (Gidlöf et al., 2021; Rebollo *et al*., 2021; Stockburger et al., 2009). Accordingly, recent studies showed that vagal stimulation induces pupil dilation and alleviates alpha oscillations, indicative of increased (visual) arousal (Collins et al., 2021; Mridha et al., 2021; Sharon et al., 2021). Relatedly, the highest increase in correlation was found in the frontal poles, which are part of the salience and default mode network, and are involved in goal-directed behavior, behavioral control and metacognition (Henri-Bhargava et al., 2018; Liu et al., 2013; Mansouri et al., 2015; Moayedi et al., 2015; Orr et al., 2015; Riedl et al., 2016) as well as in reward evaluation, planning, and decision making (Rushworth et al., 2012; Rushworth et al., 2011). This supports the idea of a relationship between hunger-dependent alterations in stomach-brain coupling and food-seeking behavior. We conclude that vagal afferent stimulation facilitates the integration of metabolic information from unimodal representations into a transmodally-extended gastric network, which may help coordinate behavioral adaptation to ensure long-term energy homeostasis.

### Gastric signaling as a common reference point for intero- and exteroception

According to Rebollo *et al*. (2021), the constant rhythmical input from the stomach may act as a common reference point for the alignment and coordination of both intero- and exteroceptive brain networks. Our finding of increased coupling in transmodal cortical regions during taVNS (vs. the predominant involvement of unimodal cortical regions during baseline) provides the first causal support for this idea. As we emulated interoceptive signals by afferently stimulating the vagus nerve, the stronger involvement of higher-order transmodal regions, which integrate activity across sensory-motor/unimodal regions (Mesulam, 1998), might mirror the coordinated hierarchical integration of altered visceral inputs across brain networks. Our results therefore suggest that the baseline gastric network reflects the continuously ongoing integration of low-level interoceptive information. As increased vagal afferent input is provided, the gastric network expands to reflect heightened integration across modalities, leading to changes in the awareness of the metabolic state of the body which may facilitate deliberate adaptations of goal-directed behavior, such as food-seeking. Crucially, the higher-order integration of the transmodal cortex during taVNS is in line with a boost in parasympathetic activity due to increased vagal signaling, whereas the baseline gastric network seems to underlie sympathetic regulation (Fedorova et al., 2020; Levinthal and Strick, 2020; Rebollo and Tallon-Baudry, 2021; Rebollo *et al*., 2021).

Strikingly, several cortical regions also show increased variance in stomach-brain coupling during taVNS (vs. sham), such as the bilateral insular cortices and the ACC. The insula displays a viscerotopic organization (Cechetto and Saper, 1987) and has not only been linked to interoception, but also saliency (Menon and Uddin, 2010). Importantly, the insula shows rich connections to several unimodal areas and processes multisensory information, including both intero- and exteroceptive signaling (Evrard, 2019). Further, the insula is involved in the parasympathetic regulation of the stomach (Levinthal and Strick, 2020). The variability of the changes in insula-stomach coupling during taVNS across our participants might reflect individual differences in interoceptive perception and processing, which seem to be associated with differences in emotional traits, genetic and child-feeding influences, psychiatric disorders, diet, and body weight (Ludwick-Rosenthal and Neufeld, 1985; Stevenson et al., 2015). Here, we did not find significant correlations between BMI or changes in gastric frequency within the regions that showed an increased variance during taVNS. Notably, we found strong associations between PLV-changes and taVNS strength in the bilateral insula (right: *r* = .587, left: *r* = .584) and the right supramarginal gyrus (*r* = .574), indicating that individual differences in stimulation strength might explain substantial variance across individuals in these regions. These findings pose the intriguing question whether stomach-brain coupling could be parametrically modulated by adjusting the stimulation strength, which requires extended designs in the future. We conclude that taVNS boosts the integration of interoceptive signaling into the transmodal cortex, modulating decision-making and arousal in a parasympathetic cortical network.

### Limitations

Despite the novel insights provided by our unique investigation combining taVNS with concurrent fMRI and EGG recordings, there are limitations that will need to be addressed in future work. First, it is not known whether the durations of taVNS alters effects on stomach-brain coupling. Our design used a duration of 10 min for each experimental phase (baseline vs. stimulation) as it is comparable to the baseline measurements of stomach-brain coupling by Rebollo *et al*. (2018). However, longer taVNS periods might elicit different changes in stomach-brain coupling as they may evoke effects on gastric frequency and motility which have mostly been demonstrated after longer stimulation periods (Steidel *et al*., 2021; Teckentrup *et al*., 2020; Zhang et al., 2021). Second, it is not known whether left-vs. right-sided stimulation leads to comparable effects on the gastric network, as lateralization differences between visceral fibers of the vagus nerve might play a role (Wang et al., 2021). Notably, the identified NTS-midbrain pathway resembles gut-induced reward after invasive stimulation of the right nodose ganglion in rodents (Han *et al*., 2018). Third, we invited participants in a metabolic state in between hunger and fullness. Future studies should systematically investigate different metabolic states and assess more extensive interoceptive measures (such as interoceptive awareness and accuracy (Legrand et al., 2021)) to further characterize the link between interoception and the gastric network. Fourth, potential interactions of the gastric network with exteroceptive input (e.g., visual or olfactory food cues) should be examined to gain insights into the alleged link between the gastric network and food-seeking behavior.

### Summary and conclusion

Procuring sufficient energy for survival is crucial and it has been hypothesized that the recently discovered gastric network plays a vital role in ensuring long-term energy homeostasis. In support of this idea, we showed that non-invasive stimulation of vagal afferents at the right ear via taVNS increases stomach-brain coupling in the NTS and the dopaminergic midbrain, in accordance with the previously established pathway for gut-induced reward (Han *et al*., 2018). Furthermore, we demonstrated increased stomach-brain coupling in transmodal cortical regions, including an increased correlation with taVNS-induced changes in subjective ratings of hunger and satiety. We conclude that taVNS is an effective method to modulate stomach-brain coupling, and an easy-to-apply, robust approach to causally study the functional contribution of the gastric network to energy metabolism in people, including patients suffering from disorders that are characterized by alterations in energy metabolism and digestion. Consequently, taVNS may open an avenue to future treatments of gastrointestinal, or more broadly, somatic symptoms of a broad array of disorders.

## STAR-Methods

### Participants

We recruited 45 healthy participants for this study. Three participants left the study by request before both sessions had been completed. One participant was omitted from this analysis because the stimulation failed to start precisely at the beginning of the stimulation phase of the resting-state fMRI (rs-fMRI) measurement. One participant was excluded due to excessive motion (> 50% of the total number of volumes exceeding a framewise displacement of 0.5 mm) in the resting-state fMRI measurement. Nine participants were excluded from analyses as they did not pass rigorous quality control of EGG and fMRI measurements, as described by Rebollo *et al*. (2018) and Wolpert et al. (2020). This yielded the final sample of 31 healthy participants (20 women, M_BMI_ = 23.42 kg/m^2^ ± 2.79 [18.79 – 30.4], M_age_= 25.39 years ± 6.58 [18 – 45]). All participants completed a telephone screening before participation to ensure eligibility. The following criteria had to be fulfilled: 1) 18-50 years of age; 2) BMI range of 18.5-30 kg/m^2^; 3) no lifetime history of brain injury, cardiovascular diseases, schizophrenia, bipolar disorder, epilepsy, diabetes, or asthma; 4) no implants (e.g. pacemaker, cochlear implant, cerebral shunt; except dental prostheses); 5) within the last 12 months: no severe substance abuse disorder, anxiety disorder (except specific phobia), obsessive-compulsive disorder, trauma- and stressor-related disorder, somatic symptom disorder or eating disorder; 6) no open wounds or impaired skin at electrode site; 7) not pregnant or nursing; 8) eligibility for MR research (i.e., no non-removable metal parts, such as piercings, no tattoos above the neck or larger than 14 cm, no claustrophobia, noise tolerability) (Teckentrup *et al*., 2021). The study was approved by the ethics committee of the University of Tübingen (reference number 235/2017BO1) and was conducted in accordance with the Declaration of Helsinki. Participants provided written informed consent at the beginning of the first session and received either monetary compensation (56€) or course credit for complete participation. Furthermore, they received additional money, office supplies, and snacks depending on their performance in two additional tasks. The full protocol is described in Teckentrup *et al*. (2021).

### Experimental procedure

Each participant completed two sessions of the same standardized protocol (randomized cross-over design, Fig. 1A). Participants received taVNS in one session, and sham stimulation in the other session with the order being determined in advance (randperm as implemented in Matlab 2018a). Participants were asked to enter the experimental session neither hungry nor full, but to refrain from consuming food or caloric beverages 1 h prior to each session. They were further asked to eat approximately 1.5 h prior to the beginning of each session to ensure comparable delays to the last meal. Next, we measured physiological parameters (heart rate, weight, waist and hip circumference) and assessed information about diet, last food, and last drink intake. Afterward, we acquired ratings of hunger, fullness, thirst, and mood based on items of the PANAS (Watson et al., 1988) using visual analog scales (VAS) on a computer.

Thereafter, participants were positioned in the fMRI scanner and all electrodes for electrogastrography (EGG) and electrocardiography (ECG) were placed according to Rebollo et al. (2018). Next, the stimulation electrode was placed at the right ear (cymba conchae for taVNS, earlobe for sham) and secured with surgical tape. We set the stimulation strength individually using VAS for pain ratings and initialized the stimulation with 1 mA and increased stepwise in 0.1 – 0.5 mA increments until the strength matched the participants’ sensation of “mild pricking” (M_taVNS_strength_ = 3.56 mA ± 1.07 [1.5 – 5]). Then, the stimulation was turned off again.

After that, we started the EGG, ECG, and fMRI measurements. We measured ECG (3 channels) and EGG (4 channels) using Brainvision Recorder (BrainProducts, Germany). First, we did an anatomical scan, followed by field maps (∼15 min) while participants completed task training. This was followed by 10 min. of baseline (no stimulation) rs-fMRI measurement. Then, the stimulation (taVNS or sham) was turned on and we measured for another ∼10 min. During rs-fMRI we showed Inscapes videos without audio to improve compliance while minimizing cognitive load and motion (Vanderwal et al., 2015). After the onset of the stimulation, taVNS or sham stimulation remained active for the rest of the session.

After rs-fMRI measurements, participants completed tasks (∼55 min). After exiting the MR scanner, participants completed the ratings of hunger, fullness, thirst, and mood again. At the end of each session, we asked participants about their belief whether they had received taVNS or sham stimulation in this session (61.3% identified the condition correctly in session 1, *p*_binomial_ = .075; 70.1% in session 2, *p*_binomial_ = .005). The second session followed the same procedure and was usually conducted 1-7 days later at the same time of the day.

### taVNS device

To administer the auricular stimulation of the vagus nerve, we used the NEMOS® stimulation device (cerbomed GmbH, Erlangen, Germany). This device follows a biphasic stimulation protocol with 30s on (frequency of 25 Hz), followed by 30s off stimulation. The stimulation intensity was set individually (possible range 0.1-5 mA) for each session until participants reported a mild pricking. We placed the stimulation electrode at the cymba conchae (taVNS) or the earlobe (sham) of the right ear (Ferstl *et al*., 2021; Frangos *et al*., 2015; Neuser *et al*., 2020; Teckentrup *et al*., 2020).

### EGG acquisition

The EGG data were recorded following the procedure published by Rebollo et al. (2018) using Brain Vision Recorder (Brain Products, Germany). To prevent the occurrence of phase differences due to unequal distances to the reference electrode, we used four bipolar electrodes for EGG acquisition that were connected to a BrainAmp amplifier (Brain Products, Germany). The data were recorded with a sampling rate of 5000 Hz with a low-pass filter of 1000 Hz and no high-pass filter. We recorded EGG (and ECG) continuously throughout the whole session. Triggers to mark the beginning and end of rs- and task-fMRI were sent using Psychtoolbox (http://psychtoolbox.org/).

### fMRI data acquisition and preprocessing

fMRI data was acquired on a Siemens 3 Tesla PRISMA magnetic resonance imaging scanner equipped with a 64-channel RF receiver head coil. Structural T1-weighted images were measured using an MP-RAGE sequence with 176 sagittal slices covering the whole brain, flip angle = 9°, matrix size = 256 × 256 and voxel size = 1 × 1 × 1 mm^3^. Field maps were acquired using a Siemens gradient echo field map sequence with short echo time (TE) = 5.19 ms and long TE = 7.65 ms (TE difference = 2.46 ms). rs-fMRI data (10 min. pre-stimulation baseline and 10 min. with concurrent stimulation) were acquired using a T2*-weighted echo-planar imaging (EPI) sequence with a multiband factor of 4, 68 axial slices with an interleaved slice order covering the whole brain (including brain stem), repetition time (TR) = 1.4 s, TE = 30 ms, flip angle = 65°, matrix size = 110 × 110, field of view = 220 × 220 mm^2^ and voxel size = 2 × 2 × 2 mm^3^. We further obtained data on the respiratory cycle based on the EGG recordings.

For preprocessing, each T1-weighted (T1w) volume was corrected for intensity non-uniformity using N4BiasFieldCorrection v2.1.0 (Tustison et al., 2010) and skull-stripped using antsBrainExtraction.sh v2.1.0 (using the OASIS template). Brain surfaces were reconstructed using recon-all from FreeSurfer v6.0.1 (Dale et al., 1999) [RRID:SCR_001847], and the brain mask estimated before was refined with a custom variation of the method to reconcile ANTs-derived and FreeSurfer-derived segmentations of the cortical gray-matter of Mindboggle (Klein et al., 2017) [RRID:SCR_002438]. Spatial normalization to the ICBM 152 Nonlinear Asymmetrical template version 2009c (Fonov et al., 2009) [RRID:SCR_008796] was performed through nonlinear registration with the antsRegistration tool of ANTs v2.1.0 (Avants et al., 2008) [RRID:SCR_004757], using brain-extracted versions of both T1w volume and template. Brain tissue segmentation of cerebrospinal fluid (CSF), white-matter (WM) and gray-matter (GM) was performed on the brain-extracted T1w using FAST (Zhang et al., 2001) [FSL v5.0.9, RRID:SCR_002823].

Functional data were slice-time corrected using 3dTshift from AFNI v16.2.07 (Cox, 1996) [RRID:SCR_005927] and motion corrected using MCFLIRT (Jenkinson et al., 2002) [FSL v5.0.9]. Distortion correction was performed using fieldmaps processed with FUGUE (Jenkinson, 2003) [FSL v5.0.9]. This was followed by co-registration to the corresponding T1w using boundary-based registration (Greve and Fischl, 2009) with 9 degrees of freedom, using bbregister [FreeSurfer v6.0.1]. Motion correcting transformations, field distortion correcting warp, BOLD-to-T1w transformation and T1w-to-template (MNI) warp were concatenated and applied in a single step using antsApplyTransforms [ANTs v2.1.0] based on Lanczos interpolation.

Physiological noise regressors were extracted by calculating the average signal inside the anatomically-derived CSF and WM masks across time using Nilearn. Framewise displacement (Power et al., 2014) was calculated for each functional run using the implementation of Nipype. Following the recommendation of Power *et al*. (2014), we calculated the number of volumes per run which exceed a framewise displacement threshold of 0.5 mm. If more than 50% of the total number of volumes exceed this threshold or less than 5 minutes of data below this threshold remain, the respective subject was excluded from further analyses.

Respiratory cycle data from the EGG electrodes was preprocessed using BrainVision Analyzer (Brain Products, Germany), FieldTrip (Oostenveld et al., 2011) and the PhysIO toolbox (Kasper et al., 2017). In brief, the EGG recordings were read into BrainVision Analyzer and the Scanner Artifact Correction was applied to remove gradient artifacts from the data before submitting exported time series to FieldTrip. As the typical respiratory rate in humans is around 0.3 Hz, the data were downsampled to 50 Hz and bandpass-filtered between 0.1 and 0.6 Hz. The respiratory time series were then read into the PhysIO toolbox and respiratory phase and respiratory volume per time were calculated. By convolution of the respiratory volume per time with the respiration response function (Birn et al., 2008), the toolbox then generated a nuisance regressor for noise correction.

## Data analysis

### EGG data

The EGG data was preprocessed using custom scripts implemented in Matlab 2019 and 2020 adapted from a pipeline published by Rebollo et al. (2018) which is available on GitHub (https://github.com/irebollo/stomach_brain_Scripts). First, the data was read out and downsampled to 10 Hz using the Fieldtrip toolbox for Matlab (Oostenveld *et al*., 2011). Next, we detrended and demeaned the signal before correcting spike and drift artifacts. Then, we separated our recordings corresponding to the performed tasks. For our analyses presented here, we only used the resting-state data, divided into baseline and stimulation phases. We obtained the frequency spectrum of the stomach corresponding to the range between 0.03-0.07 Hz by performing a fast Fourier transform. Previous studies suggest that the gastric frequency is reliably centered around 0.05 Hz in healthy humans (Wolpert *et al*., 2020), so we used a Cauchy distribution centered around 0.05 Hz to assign peaks close to the expected frequency a weigher weight a priori to improve robustness of peak detection. Then, we conducted a visual quality control to determine the usability of the data. Our criteria for this were a clearly visible peak in the expected frequency spectrum (0.03-0.07 Hz) with a power ≥15 µV which was congruent across channels and identifiable for both phases (baseline and stimulation). Next, we defined the gastric frequency of each participant based on the strongest usable EGG channel. We prioritized the highest power during the stimulation phase, as long as the same channel was usable in both phases.

Afterward, we bandpass-filtered the EGG time series for both phases (baseline and stimulation) around the respective center frequency. We then downsampled the EGG time series according to the fMRI sampling rate (0.714 Hz). Thereafter, we performed a Hilbert transformation to obtain the phase information of the signal.

### fMRI data

After preprocessing, we extracted the BOLD time series from the rs-fMRI measurements for each voxel inside the brain mask provided by SPM. We then normalized and z-scored these time series. We denoised the signal using regressors for movement, cerebrospinal fluid, white matter, and respiration using the TAPAS toolbox (Frässle et al., 2021). Then, we split the time series into the same experimental phases as the EGG data (baseline and stimulation) and bandpass-filtered both parts around the corresponding individual EGG peak frequency. Next, we performed a Hilbert transformation to obtain the phase information of the BOLD time series.

### Phase coupling

To determine the phase coupling between the EGG and BOLD time series, we computed the phase-locking value (PLV) for each voxel (Rebollo *et al*., 2018). The PLV describes the synchrony between two signals, for example, that they de-/accelerate simultaneously, and can take values from 0 (no synchrony) to 1 (perfect synchrony) (Fig. 1A).

To test whether observed phase coupling exceeds levels occurring by chance, we used a permutation approach. We generated permuted phase sequences by computing chance-PLV (CPLV), by circularly shifting the EGG phase signal (signal was cut from the end and prepended the beginning) and computing the PLV for each shift. This procedure disassociates the temporal structure between the EGG and the BOLD time series, effectively removing meaningful synchrony between the two signals. The EGG signal was shifted by at least one minute to ensure sufficient discrepancy to the empirical PLV. Consequently, time series were shifted 343 times, resulting in 343 shifted PLV per voxel. Each voxel’s CPLV was then defined as the mean of all shifted PLV of this specific voxel (Rebollo *et al*., 2018).

To replicate the resting-state stomach-brain network as previously published by Rebollo *et al*. (2018), we first analyzed data from the baseline phase only using the same protocol published in Rebollo *et al*. (2018). We compared each voxel’s empirical PLV to the CPLV using a paired *t*-test and thresholded the resulting maps on the cluster level, determining significant clusters using a Monte-Carlo permutation test (cluster-α = 0.005, α = 0.025, 10,000 permutations).

Next, we analyzed the effect of taVNS on stomach-brain coupling using full-factorical models in SPM 12 and Matlab by computing the interaction effect of Stimulation [taVNS versus sham] × Time [baseline versus stimulation] and added the order of the stimulation conditions as a covariate. Moreover, we analyzed whether taVNS increases the variance in PLV changes using *F*-tests (taVNS divided by sham variance). We performed an exploratory analysis of the correlation between PLV changes (mean over region) and changes in gastric myoelectric frequency, BMI, and taVNS strength to evaluate potential influential factors that could explain part of the variance. To investigate if metabolic state mediates this effect of taVNS on stomach-brain coupling, we calculated the correlation between changes in coupling due to taVNS (ΔPLV_taVNS_ [stimulation-baseline] - ΔPLV_sham_ -[stimulation-baseline]) and changes in metabolic state over the session (post [hunger-satiety] - pre [hunger-satiety]).

To analyze how taVNS affects the coupling of functional networks, we assigned each brain region within the extended Harvard-Oxford atlas to one of the 7 cortical resting-state networks as as published by Yeo et al. (2011) (DMN, ventral and dorsal attention, visual, somatomotor, frontoparietal, limbic) or a subcortical anatomical network (brainstem, midbrain, diencephalon). The assignment was based on the largest overlap between each region defined within the extended Harvard-Oxford atlas and the network maps provided by Yeo et al. (2011). To evaluate which functional networks show the most substantial change after taVNS, we used the t-maps resulting from the full factorial models described above and calculated voxel-wise Cohen’s d_z_ as a measure of effect size by dividing the *t*-values by the square-root of the sample size (n = 31). Then, we averaged *d*_*z*_ across all voxels assigned to the same network. To assess how the voxels affected by taVNS are positioned in the cortical hierarchy, we assigned each voxel within the t-maps of the SPM-models for Stimulation × Time and the correlation with the metabolic state (details described above, maps available on NeuroVault) to one of the first two cortical resting-state gradients (gradient 1: unimodal to transmodal, gradient 2: somatosensory/somatomotor to visual) as published by Margulies *et al*. (2016). We then performed two-sample Kolmogorov-Smirnov tests (MATLAB function kstest2) between the standard cortical gradients and the respective distributions during baseline and stimulation to evaluate how taVNS changes large-scale connectivity patterns.

### Metabolic state

To analyze the assessed subjective metabolic state (VAS before and after measurement), we used a linear mixed effects model using the lme4 package v1.1-21 in R (Bates et al., 2015): *rating ∼ stimulation * time+ (1 + stimulation* | *ID)* (BIC = 1292.5).

### Statistical threshold and software

Our analyses were conducted with Matlab v2019 & v2020 and SPM12. For preprocessing of the EGG data, we used the Fieldtrip toolbox for Matlab (Oostenveld *et al*., 2011). The fMRI data were preprocessed using the standardized FMRIPREP pipeline (https://github.com/poldracklab/fmriprep) v20.1.1 (Esteban et al., 2019) based on Nipype (Gorgolewski et al., 2011) [RRID:SCR_002502] and Nilearn (Abraham et al., 2014) [RRID:SCR_001362]. For parcellation of brain maps, we used an extended version of the Harvard-Oxford atlas (Desikan et al., 2006), which includes the Reinforcement Learning Atlas (https://osf.io/jkzwp/) for extended coverage of subcortical nuclei and the AAL cerebellum ROIs (Tzourio-Mazoyer et al., 2002). We used RStudio v1.2.5 with the ggplot2 package v3.3.2 (Wickham, 2016), including the tidyverse package v1.3, the viridis color package v0.5.1 (Garnier, 2021), the gghighlight package v0.3.2 (https://github.com/yutannihilation/gghighlight/), and the cowplot package v1.1.0 (https://wilkelab.org/cowplot/) for plotting. We considered α < 0.05 as significant, except for FWE cluster-corrected t-maps, where we considered *p* < .001 (*k* ≥ 40) as significant. The cluster extent threshold for the SPM analyses was estimated with AlphaSim (Song et al., 2011) based on the smoothness of the maps (α_corrected_ < .05, 2,000 iterations). For the baseline results, we thresholded the PLV vs CPLV maps using a Monte-Carlo permutation test (cluster-α = 0.005, α = 0.025, 10,000 permutations).

## Data and code availability

The analyses are based on code published by Rebollo *et al*. (2018) that is available on: https://github.com/irebollo/stomach_brain_Scripts. All unthresholded group-level maps of the results are uploaded on NeuroVault: https://neurovault.org/collections/QNGZBQGF/.

## Acknowledgment

We thank Sandra Neubert, Vinzent Wolf, Franziska Müller, Corinna Schulz, Wy Ming Lin, Monja Neuser and Franziska Kräutlein for help with data acquisition. The study was supported by the University of Tübingen, Faculty of Medicine fortune grant #2453-0-0 and a postdoctoral scholarship 32-04/19 provided by the Daimler and Benz Foundation awarded to NBK.

## Author contributions

NBK was responsible for the study concept and design. VT & SM collected data under supervision by NBK. NBK conceived the method and SM & VT processed the data. SM & VT performed the data analysis and NBK contributed to analyses. SM, VT & NBK wrote the manuscript. All authors contributed to the interpretation of findings, provided critical revision of the manuscript for important intellectual content, and approved the final version for publication.

## References

Abraham, A., Pedregosa, F., Eickenberg, M., Gervais, P., Mueller, A., Kossaifi, J., Gramfort, A., Thirion, B., and Varoquaux, G. (2014). Machine learning for neuroimaging with scikit-learn. Front Neuroinform 8, 14. 10.3389/fninf.2014.00014.

Alhadeff, A.L., and Grill, H.J. (2014). Hindbrain nucleus tractus solitarius glucagon-like peptide-1 receptor signaling reduces appetitive and motivational aspects of feeding. American Journal of Physiology-Regulatory, Integrative and Comparative Physiology 307, R465–R470. 10.1152/ajpregu.00179.2014.

Allen, M., Levy, A., Parr, T., and Friston, K.J. (2019). In the Body’s Eye: The Computational Anatomy of Interoceptive Inference. bioRxiv, 603928. 10.1101/603928.

Anselmi, L., Toti, L., Bove, C., Hampton, J., and Travagli, R.A. (2017). A Nigro-Vagal Pathway Controls Gastric Motility and Is Affected in a Rat Model of Parkinsonism. Gastroenterology 153, 1581–1593. 10.1053/j.gastro.2017.08.069.

Aurora, S.K., Shrewsbury, S.B., Ray, S., Hindiyeh, N., and Nguyen, L. (2021). A link between gastrointestinal disorders and migraine: Insights into the gut-brain connection. Headache. 10.1111/head.14099.

Avants, B.B., Epstein, C.L., Grossman, M., and Gee, J.C. (2008). Symmetric diffeomorphic image registration with cross-correlation: evaluating automated labeling of elderly and neurodegenerative brain. Med Image Anal 12, 26–41. 10.1016/j.media.2007.06.004.

Azzalini, D., Rebollo, I., and Tallon-Baudry, C. (2019). Visceral Signals Shape Brain Dynamics and Cognition. Trends Cogn Sci 23, 488–509. 10.1016/j.tics.2019.03.007.

Bates, D., Mächler, M., Bolker, B., and Walker, S. (2015). Fitting Linear Mixed-Effects Models Using lme4. Journal of Statistical Software; Vol 1, Issue 1 (2015).

Berthoud, H.R., and Neuhuber, W.L. (2000). Functional and chemical anatomy of the afferent vagal system. Auton Neurosci 85, 1–17. 10.1016/S1566-0702(00)00215-0.

Birn, R.M., Smith, M.A., Jones, T.B., and Bandettini, P.A. (2008). The respiration response function: the temporal dynamics of fMRI signal fluctuations related to changes in respiration. Neuroimage 40, 644–654. 10.1016/j.neuroimage.2007.11.059.

Boughter, J.D., Jr., Lu, L., Saites, L.N., and Tokita, K. (2019). Sweet and bitter taste stimuli activate VTA projection neurons in the parabrachial nucleus. Brain Res 1714, 99–110. 10.1016/j.brainres.2019.02.027.

Breen, D.P., Halliday, G.M., and Lang, A.E. (2019). Gut-brain axis and the spread of alphasynuclein pathology: Vagal highway or dead end? Mov Disord 34, 307–316. 10.1002/mds.27556.

Cechetto, D.F., and Saper, C.B. (1987). Evidence for a viscerotopic sensory representation in the cortex and thalamus in the rat. J Comp Neurol 262, 27–45. 10.1002/cne.902620104.

Choe, A.S., Tang, B., Smith, K.R., Honari, H., Lindquist, M.A., Caffo, B.S., and Pekar, J.J. (2021). Phase-locking of resting-state brain networks with the gastric basal electrical rhythm. PLoS One 16, e0244756. 10.1371/journal.pone.0244756.

Collins, L., Boddington, L., Steffan, P.J., and McCormick, D. (2021). Vagus nerve stimulation induces widespread cortical and behavioral activation. Curr Biol 31, 2088–2098 e2083. 10.1016/j.cub.2021.02.049.

Cox, R.W. (1996). AFNI: software for analysis and visualization of functional magnetic resonance neuroimages. Comput Biomed Res 29, 162–173. 10.1006/cbmr.1996.0014.

Dale, A.M., Fischl, B., and Sereno, M.I. (1999). Cortical surface-based analysis. I. Segmentation and surface reconstruction. Neuroimage 9, 179–194. 10.1006/nimg.1998.0395.

Desikan, R.S., Segonne, F., Fischl, B., Quinn, B.T., Dickerson, B.C., Blacker, D., Buckner, R.L., Dale, A.M., Maguire, R.P., Hyman, B.T., et al. (2006). An automated labeling system for subdividing the human cerebral cortex on MRI scans into gyral based regions of interest. Neuroimage 31, 968–980. 10.1016/j.neuroimage.2006.01.021.

Dunckley, P., Wise, R.G., Fairhurst, M., Hobden, P., Aziz, Q., Chang, L., and Tracey, I. (2005). A comparison of visceral and somatic pain processing in the human brainstem using functional magnetic resonance imaging. J Neurosci 25, 7333–7341. 10.1523/JNEUROSCI.1100-05.2005.

Esteban, O., Markiewicz, C.J., Blair, R.W., Moodie, C.A., Isik, A.I., Erramuzpe, A., Kent, J.D., Goncalves, M., DuPre, E., Snyder, M., et al. (2019). fMRIPrep: a robust preprocessing pipeline for functional MRI. Nat Methods 16, 111–116. 10.1038/s41592-018-0235-4.

Evrard, H.C. (2019). The Organization of the Primate Insular Cortex. Front Neuroanat 13, 43. 10.3389/fnana.2019.00043.

Fedorova, T.D., Knudsen, K., Hartmann, B., Holst, J.J., Viborg Mortensen, F., Krogh, K., and Borghammer, P. (2020). In vivo positron emission tomography imaging of decreased parasympathetic innervation in the gut of vagotomized patients. Neurogastroenterol Motil 32, e13759. 10.1111/nmo.13759.

Fernandes, A.B., Alves da Silva, J., Almeida, J., Cui, G., Gerfen, C. R., Costa, R.M., and Oliveira-Maia, A.J. (2020). Postingestive Modulation of Food Seeking Depends on Vagus-Mediated Dopamine Neuron Activity. Neuron 106, 778-788.e776. 10.1016/j.neuron.2020.03.009.

Ferstl, M., Teckentrup, V., Lin, W.M., Krautlein, F., Kuhnel, A., Klaus, J., Walter, M., and Kroemer, N.B. (2021). Non-invasive vagus nerve stimulation boosts mood recovery after effort exertion. Psychol Med, 1-11. 10.1017/S0033291720005073.

Folgueira, C., Seoane, L.M., and Casanueva, F.F. (2014). The brain-stomach connection. Front Horm Res 42, 83–92. 10.1159/000358316.

Fonov, V.S., Evans, A.C., McKinstry, R.C., Almli, C.R., and Collins, D.L. (2009). Unbiased nonlinear average age-appropriate brain templates from birth to adulthood. NeuroImage 47, S102. https://doi.org/10.1016/S1053-8119(09)70884-5.

Frangos, E., Ellrich, J., and Komisaruk, B.R. (2015). Non-invasive Access to the Vagus Nerve Central Projections via Electrical Stimulation of the External Ear: fMRI Evidence in Humans. Brain Stimul 8, 624–636. 10.1016/j.brs.2014.11.018.

Frässle, S., Aponte, E.A., Bollmann, S., Brodersen, K.H., Do, C.T., Harrison, O.K., Harrison, S.J., Heinzle, J., Iglesias, S., Kasper, L., et al. (2021). TAPAS: An Open-Source Software Package for Translational Neuromodeling and Computational Psychiatry. Frontiers in Psychiatry 12. 10.3389/fpsyt.2021.680811.

Garnier, S., Ross, Noam, Rudis, Robert, Camargo, Pedro A, Sciaini, Marco, Scherer, Cédric (2021). viridis - Colorblind-Friendly Color Maps for R.

Gidlöf, K., Ares, G., Aschemann-Witzel, J., and Otterbring, T. (2021). Give us today our daily bread: The effect of hunger on consumers’ visual attention towards bread and the role of time orientation. Food Quality and Preference 88, 104079.

Giraudier, M., Ventura-Bort, C., and Weymar, M. (2020). Transcutaneous Vagus Nerve Stimulation (tVNS) Improves High-Confidence Recognition Memory but Not Emotional Word Processing. Frontiers in Psychology 11. 10.3389/fpsyg.2020.01276.

Gorgolewski, K., Burns, C.D., Madison, C., Clark, D., Halchenko, Y.O., Waskom, M.L., and Ghosh, S.S. (2011). Nipype: a flexible, lightweight and extensible neuroimaging data processing framework in python. Front Neuroinform 5, 13. 10.3389/fninf.2011.00013.

Greve, D.N., and Fischl, B. (2009). Accurate and robust brain image alignment using boundary-based registration. Neuroimage 48, 63–72. 10.1016/j.neuroimage.2009.06.060.

Han, W., Tellez, L.A., Perkins, M.H., Perez, I.O., Qu, T., Ferreira, J., Ferreira, T.L., Quinn, D., Liu, Z.W., Gao, X.B., et al. (2018). A Neural Circuit for Gut-Induced Reward. Cell 175, 887–888. 10.1016/j.cell.2018.10.018.

He, W., Jing, X.H., Zhu, B., Zhu, X.L., Li, L., Bai, W.Z., and Ben, H. (2013). The auriculo-vagal afferent pathway and its role in seizure suppression in rats. BMC Neurosci 14, 85. 10.1186/1471-2202-14-85.

Heimrich, K.G., Jacob, V.Y.P., Schaller, D., Stallmach, A., Witte, O.W., and Prell, T. (2019). Gastric dysmotility in Parkinson’s disease is not caused by alterations of the gastric pacemaker cells. npj Parkinson’s Disease 5. 10.1038/s41531-019-0087-3.

Henri-Bhargava, A., Stuss, D.T., and Freedman, M. (2018). Clinical Assessment of Prefrontal Lobe Functions. Continuum (Minneap Minn) 24, 704–726. 10.1212/CON.0000000000000609.

Holtmann, G., and Talley, N.J. (2014). The stomach-brain axis. Best Pract Res Clin Gastroenterol 28, 967–979. 10.1016/j.bpg.2014.10.001.

Hong, G.S., Pintea, B., Lingohr, P., Coch, C., Randau, T., Schaefer, N., Wehner, S., Kalff, J.C., and Pantelis, D. (2019). Effect of transcutaneous vagus nerve stimulation on muscle activity in the gastrointestinal tract (transVaGa): a prospective clinical trial. Int J Colorectal Dis 34, 417-422 10.1007/s00384-018-3204-6.

Huang, M.H., Wang, Y.P., Wu, P.S., Chan, Y.E., Cheng, C.M., Yang, C.H., Tsai, S.J., Lu, C.L., and Tsai, C.F. (2021). Association between gastrointestinal symptoms and depression among older adults in Taiwan: A cross-sectional study. J Chin Med Assoc 84, 331–335. 10.1097/JCMA.0000000000000460.

Hussain, M.M., and Pan, X. (2009). Clock genes, intestinal transport and plasma lipid homeostasis. Trends in Endocrinology & Metabolism 20, 177–185. 10.1016/j.tem.2009.01.001.

Jacobs, H.I., Riphagen, J.M., Razat, C.M., Wiese, S., and Sack, A.T. (2015). Transcutaneous vagus nerve stimulation boosts associative memory in older individuals. Neurobiol Aging 36, 1860–1867. 10.1016/j.neurobiolaging.2015.02.023.

Jenkinson, M. (2003). Fast, automated, N-dimensional phase-unwrapping algorithm. Magn Reson Med 49, 193–197. 10.1002/mrm.10354.

Jenkinson, M., Bannister, P., Brady, M., and Smith, S. (2002). Improved optimization for the robust and accurate linear registration and motion correction of brain images. Neuroimage 17, 825–841. 10.1016/s1053-8119(02)91132-8.

Kanoski, S.E., Alhadeff, A.L., Fortin, S.M., Gilbert, J.R., and Grill, H.J. (2013). Leptin Signaling in the Medial Nucleus Tractus Solitarius Reduces Food Seeking and Willingness to Work for Food. Neuropsychopharmacology 39, 605–613. 10.1038/npp.2013.235.

Kasper, L., Bollmann, S., Diaconescu, A.O., Hutton, C., Heinzle, J., Iglesias, S., Hauser, T.U., Sebold, M., Manjaly, Z.M., Pruessmann, K.P., and Stephan, K.E. (2017). The PhysIO Toolbox for Modeling Physiological Noise in fMRI Data. J Neurosci Methods 276, 56–72. 10.1016/j.jneumeth.2016.10.019.

Kaut, O., Janocha, L., Weismuller, T.J., and Wullner, U. (2019). Transcutaneous vagal nerve stimulation improves gastroenteric complaints in Parkinson’s disease patients. NeuroRehabilitation 45, 449–451. 10.3233/NRE-192909.

Klein, A., Ghosh, S.S., Bao, F.S., Giard, J., Hame, Y., Stavsky, E., Lee, N., Rossa, B., Reuter, M., Chaibub Neto, E., and Keshavan, A. (2017). Mindboggling morphometry of human brains. PLoS Comput Biol 13, e1005350. 10.1371/journal.pcbi.1005350.

Kraus, T., Hösl, K., Kiess, O., Schanze, A., Kornhuber, J., and Forster, C. (2007). BOLD fMRI deactivation of limbic and temporal brain structures and mood enhancing effect by transcutaneous vagus nerve stimulation. Journal of Neural Transmission 114, 1485–1493. 10.1007/s00702-007-0755-z.

Legrand, N., Nikolova, N., Correa, C., Brændholt, M., Stuckert, A., Kildahl, N., Vejlø, M., Fardo, F., and Allen, M. (2021). The heart rate discrimination task: a psychophysical method to estimate the accuracy and precision of interoceptive beliefs. bioRxiv.

Levakov, G., Kaplan, A., Yaskolka Meir, A., Rinott, E., Tsaban, G., Zelicha, H., Meiran, N., Shelef, I., Shai, I., and Avidan, G. (2021). Neural correlates of future weight loss reveal a possible role for brain-gastric interactions. Neuroimage 224, 117403. 10.1016/j.neuroimage.2020.117403.

Levinthal, D.J., and Strick, P.L. (2020). Multiple areas of the cerebral cortex influence the stomach. Proc Natl Acad Sci U S A 117, 13078–13083. 10.1073/pnas.2002737117.

Liu, H., Qin, W., Li, W., Fan, L., Wang, J., Jiang, T., and Yu, C. (2013). Connectivity-based parcellation of the human frontal pole with diffusion tensor imaging. J Neurosci 33, 6782–6790. 10.1523/JNEUROSCI.4882-12.2013.

Lubomski, M., Davis, R.L., and Sue, C.M. (2020). Gastrointestinal dysfunction in Parkinson’s disease. J Neurol 267, 1377–1388. 10.1007/s00415-020-09723-5.

Ludwick-Rosenthal, R., and Neufeld, R.W. (1985). Heart beat interoception: a study of individual differences. Int J Psychophysiol 3, 57–65. 10.1016/0167-8760(85)90020-0.

Lundgren, O. (1983). Vagal control of the motor functions of the lower esophageal sphincter and the stomach. J Auton Nerv Syst 9, 185–197. 10.1016/0165-1838(83)90140-6.

Mandal, A., Prabhavalkar, K.S., and Bhatt, L.K. (2018). Gastrointestinal hormones in regulation of memory. Peptides 102, 16–25. 10.1016/j.peptides.2018.02.003.

Mansouri, F.A., Buckley, M.J., Mahboubi, M., and Tanaka, K. (2015). Behavioral consequences of selective damage to frontal pole and posterior cingulate cortices. Proc Natl Acad Sci U S A 112, E3940–3949. 10.1073/pnas.1422629112.

Margulies, D.S., Ghosh, S.S., Goulas, A., Falkiewicz, M., Huntenburg, J.M., Langs, G., Bezgin, G., Eickhoff, S.B., Castellanos, F.X., Petrides, M., et al. (2016). Situating the default-mode network along a principal gradient of macroscale cortical organization. Proc Natl Acad Sci U S A 113, 12574–12579. 10.1073/pnas.1608282113.

Marrinan, S., Emmanuel, A.V., and Burn, D.J. (2014). Delayed gastric emptying in Parkinson’s disease. Mov Disord 29, 23–32. 10.1002/mds.25708.

Mattes, R.D., Hunter, S.R., and Higgins, K.A. (2019). Sensory, gastric, and enteroendocrine effects of carbohydrates, fat, and protein on appetite. Current Opinion in Endocrine and Metabolic Research 4, 14–20. 10.1016/j.coemr.2018.09.002.

Mayer, E.A. (2011). Gut feelings: the emerging biology of gut–brain communication. Nature Reviews Neuroscience 12, 453–466. 10.1038/nrn3071.

Mazzoni, P., Hristova, A., and Krakauer, J.W. (2007). Why Don’t We Move Faster? Parkinson’s Disease, Movement Vigor, and Implicit Motivation. Journal of Neuroscience 27, 7105–7116. 10.1523/jneurosci.0264-07.2007.

Menon, V., and Uddin, L.Q. (2010). Saliency, switching, attention and control: a network model of insula function. Brain Struct Funct 214, 655–667. 10.1007/s00429-010-0262-0.

Mesulam, M.M. (1998). From sensation to cognition. Brain 121 (Pt 6), 1013–1052. 10.1093/brain/121.6.1013.

Moayedi, M., Salomons, T.V., Dunlop, K.A., Downar, J., and Davis, K.D. (2015). Connectivity-based parcellation of the human frontal polar cortex. Brain Struct Funct 220, 2603–2616. 10.1007/s00429-014-0809-6.

Mrabet, S., Ben Ali, N., Achouri, A., Dabbeche, R., Najjar, T., Haouet, S., and Belal, S. (2016). Gastrointestinal Dysfunction and Neuropathologic Correlations in Parkinson Disease. J Clin Gastroenterol 50, e85–90. 10.1097/MCG.0000000000000606.

Mridha, Z., de Gee, J.W., Shi, Y., Alkashgari, R., Williams, J., Suminski, A., Ward, M.P., Zhang, W., and McGinley, M.J. (2021). Graded recruitment of pupil-linked neuromodulation by parametric stimulation of the vagus nerve. Nat Commun 12, 1539. 10.1038/s41467-021-21730-2.

Mussell, M., Kroenke, K., Spitzer, R.L., Williams, J.B., Herzog, W., and Lowe, B. (2008). Gastrointestinal symptoms in primary care: prevalence and association with depression and anxiety. J Psychosom Res 64, 605–612. 10.1016/j.jpsychores.2008.02.019.

Neuhuber, W.L., and Berthoud, H.-R. (2021). Functional anatomy of the vagus system – Emphasis on the somato-visceral interface. Autonomic Neuroscience, 102887. https://doi.org/10.1016/j.autneu.2021.102887.

Neuser, M.P., Teckentrup, V., Kühnel, A., Hallschmid, M., Walter, M., and Kroemer, N.B. (2020). Vagus nerve stimulation boosts the drive to work for rewards. Nat Commun 11, 3555 10.1038/s41467-020-17344-9.

Nord, C.L., Dalmaijer, E.S., Armstrong, T., Baker, K., and Dalgleish, T. (2021). A Causal Role for Gastric Rhythm in Human Disgust Avoidance. Curr Biol 31, 629–634 e623. 10.1016/j.cub.2020.10.087.

Oostenveld, R., Fries, P., Maris, E., and Schoffelen, J.M. (2011). FieldTrip: Open source software for advanced analysis of MEG, EEG, and invasive electrophysiological data. Comput Intell Neurosci 2011, 156869 10.1155/2011/156869.

Orr, J.M., Smolker, H.R., and Banich, M.T. (2015). Organization of the Human Frontal Pole Revealed by Large-Scale DTI-Based Connectivity: Implications for Control of Behavior. PLoS One 10, e0124797. 10.1371/journal.pone.0124797.

Power, J.D., Mitra, A., Laumann, T.O., Snyder, A.Z., Schlaggar, B.L., and Petersen, S.E. (2014). Methods to detect, characterize, and remove motion artifact in resting state fMRI. Neuroimage 84, 320–341. 10.1016/j.neuroimage.2013.08.048.

Powley, T.L., Spaulding, R.A., and Haglof, S.A. (2011). Vagal afferent innervation of the proximal gastrointestinal tract mucosa: chemoreceptor and mechanoreceptor architecture. J Comp Neurol 519, 644–660. 10.1002/cne.22541.

Rassi, E., Dorffner, G., Gruber, W., Schabus, M., and Klimesch, W. (2019). Coupling and Decoupling between Brain and Body Oscillations. Neurosci Lett 711, 134401. 10.1016/j.neulet.2019.134401.

Rebollo, I., Devauchelle, A.D., Beranger, B., and Tallon-Baudry, C. (2018). Stomach-brain synchrony reveals a novel, delayed-connectivity resting-state network in humans. Elife 7. 10.7554/eLife.33321.

Rebollo, I., and Tallon-Baudry, C. (2021). The sensory and motor components of the cortical hierarchy are coupled to the rhythm of the stomach during rest. bioRxiv, 2021.2005.2026.445829 10.1101/2021.05.26.445829.

Rebollo, I., Wolpert, N., and Tallon-Baudry, C. (2021). Brain–stomach coupling: Anatomy, functions, and future avenues of research. Current Opinion in Biomedical Engineering 18. 10.1016/j.cobme.2021.100270.

Riedl, V., Utz, L., Castrillon, G., Grimmer, T., Rauschecker, J.P., Ploner, M., Friston, K.J., Drzezga, A., and Sorg, C. (2016). Metabolic connectivity mapping reveals effective connectivity in the resting human brain. Proc Natl Acad Sci U S A 113, 428–433. 10.1073/pnas.1513752113.

Rushworth, M.F., Kolling, N., Sallet, J., and Mars, R.B. (2012). Valuation and decision-making in frontal cortex: one or many serial or parallel systems? Curr Opin Neurobiol 22, 946–955. 10.1016/j.conb.2012.04.011.

Rushworth, M.F., Noonan, M.P., Boorman, E.D., Walton, M.E., and Behrens, T.E. (2011). Frontal cortex and reward-guided learning and decision-making. Neuron 70, 1054–1069. 10.1016/j.neuron.2011.05.014.

Santos-Garcia, D., de Deus, T., Tejera-Perez, C., Exposito-Ruiz, I., Suarez-Castro, E., Carpintero, P., and Macias-Arribi, M. (2015). [Gastroparesis and other gastrointestinal symptoms in Parkinson’s disease]. Rev Neurol 61, 261–270.

Sclocco, R., Garcia, R.G., Kettner, N.W., Isenburg, K., Fisher, H.P., Hubbard, C.S., Ay, I., Polimeni, J.R., Goldstein, J., Makris, N., et al. (2019). The influence of respiration on brainstem and cardiovagal response to auricular vagus nerve stimulation: A multimodal ultrahigh-field (7T) fMRI study. Brain Stimulation 12, 911–921. 10.1016/j.brs.2019.02.003.

Shapiro, R.E., and Miselis, R.R. (1985). The central organization of the vagus nerve innervating the stomach of the rat. J Comp Neurol 238, 473–488. 10.1002/cne.902380411.

Sharon, O., Fahoum, F., and Nir, Y. (2021). Transcutaneous Vagus Nerve Stimulation in Humans Induces Pupil Dilation and Attenuates Alpha Oscillations. J Neurosci 41, 320–330. 10.1523/JNEUROSCI.1361-20.2020.

Soderquist, F., Syk, M., Just, D., Kurbalija Novicic, Z., Rasmusson, A.J., Hellstrom, P.M., Ramklint, M., and Cunningham, J.L. (2020). A cross-sectional study of gastrointestinal symptoms, depressive symptoms and trait anxiety in young adults. BMC Psychiatry 20, 535. 10.1186/s12888-020-02940-2.

Song, X.-W., Dong, Z.-Y., Long, X.-Y., Li, S.-F., Zuo, X.-N., Zhu, C.-Z., He, Y., Yan, C.-G., and Zang, Y.-F. (2011). REST: a toolkit for resting-state functional magnetic resonance imaging data processing. PloS one 6, e25031–e25031. 10.1371/journal.pone.0025031.

Steidel, K., Krause, K., Menzler, K., Strzelczyk, A., Immisch, I., Fuest, S., Gorny, I., Mross, P., Hakel, L., Schmidt, L., et al. (2021). Transcutaneous auricular vagus nerve stimulation influences gastric motility: A randomized, double-blind trial in healthy individuals. Brain Stimul. 10.1016/j.brs.2021.06.006.

Stevenson, R.J., Mahmut, M., and Rooney, K. (2015). Individual differences in the interoceptive states of hunger, fullness and thirst. Appetite 95, 44–57. 10.1016/j.appet.2015.06.008.

Stockburger, J., Schmalzle, R., Flaisch, T., Bublatzky, F., and Schupp, H.T. (2009). The impact of hunger on food cue processing: an event-related brain potential study. Neuroimage 47, 1819–1829. 10.1016/j.neuroimage.2009.04.071.

Suarez, A.N., Hsu, T.M., Liu, C.M., Noble, E.E., Cortella, A.M., Nakamoto, E.M., Hahn, J.D., de Lartigue, G., and Kanoski, S.E. (2018). Gut vagal sensory signaling regulates hippocampus function through multi-order pathways. Nature Communications 9. 10.1038/s41467-018-04639-1.

Svensson, E., Horváth-Puhó, E., Thomsen, R.W., Djurhuus, J.C., Pedersen, L., Borghammer, P., and Sørensen, H.T. (2015). Vagotomy and subsequent risk of Parkinson’s disease. Annals of Neurology 78, 522–529. 10.1002/ana.24448.

Teckentrup, V., Krylova, M., Jamalabadi, H., Neubert, S., Neuser, M.P., Hartig, R., Fallgatter, A.J., Walter, M., and Kroemer, N.B. (2021). Brain signaling dynamics after vagus nerve stimulation. bioRxiv, 2021.2006.2028.450171. 10.1101/2021.06.28.450171.

Teckentrup, V., Neubert, S., Santiago, J.C.P., Hallschmid, M., Walter, M., and Kroemer, N.B. (2020). Non-invasive stimulation of vagal afferents reduces gastric frequency. Brain Stimul 13, 470–473. 10.1016/j.brs.2019.12.018.

Tustison, N.J., Avants, B.B., Cook, P.A., Zheng, Y., Egan, A., Yushkevich, P.A., and Gee, J.C. (2010). N4ITK: improved N3 bias correction. IEEE Trans Med Imaging 29, 1310–1320. 10.1109/TMI.2010.2046908.

Tzourio-Mazoyer, N., Landeau, B., Papathanassiou, D., Crivello, F., Etard, O., Delcroix, N., Mazoyer, B., and Joliot, M. (2002). Automated anatomical labeling of activations in SPM using a macroscopic anatomical parcellation of the MNI MRI single-subject brain. Neuroimage 15, 273–289. 10.1006/nimg.2001.0978.

Vanderwal, T., Kelly, C., Eilbott, J., Mayes, L.C., and Castellanos, F.X. (2015). Inscapes: A movie paradigm to improve compliance in functional magnetic resonance imaging. Neuroimage 122, 222–232. 10.1016/j.neuroimage.2015.07.069.

Varga, S., and Heck, D.H. (2017). Rhythms of the body, rhythms of the brain: Respiration, neural oscillations, and embodied cognition. Consciousness and Cognition 56, 77–90. 10.1016/j.concog.2017.09.008.

Vazquez-Oliver, A., Brambilla-Pisoni, C., Domingo-Gainza, M., Maldonado, R., Ivorra, A., and Ozaita, A. (2020). Auricular transcutaneous vagus nerve stimulation improves memory persistence in naive mice and in an intellectual disability mouse model. Brain Stimul 13, 494–498. 10.1016/j.brs.2019.12.024.

Waise, T.M.Z., Dranse, H.J., and Lam, T.K.T. (2018). The metabolic role of vagal afferent innervation. Nat Rev Gastroenterol Hepatol 15, 625–636. 10.1038/s41575-018-0062-1.

Wang, Y., Li, S.Y., Wang, D., Wu, M.Z., He, J.K., Zhang, J.L., Zhao, B., Hou, L.W., Wang, J.Y., Wang, L., et al. (2021). Transcutaneous Auricular Vagus Nerve Stimulation: From Concept to Application. Neurosci Bull 37, 853–862. 10.1007/s12264-020-00619-y.

Waterson, M.J., and Horvath, T.L. (2015). Neuronal Regulation of Energy Homeostasis: Beyond the Hypothalamus and Feeding. Cell Metab 22, 962–970. 10.1016/j.cmet.2015.09.026.

Watson, D., Clark, L.A., and Tellegen, A. (1988). Development and validation of brief measures of positive and negative affect: the PANAS scales. J Pers Soc Psychol 54, 1063–1070. 10.1037//0022-3514.54.6.1063.

Wickham, H. (2016). ggplot2: Elegant Graphics for Data Analysis (Springer-Verlag New York).

Wolpert, N., Rebollo, I., and Tallon-Baudry, C. (2020). Electrogastrography for psychophysiological research: Practical considerations, analysis pipeline, and normative data in a large sample. Psychophysiology 57, e13599. 10.1111/psyp.13599.

Yakunina, N., Kim, S.S., and Nam, E.-C. (2017). Optimization of Transcutaneous Vagus Nerve Stimulation Using Functional MRI. Neuromodulation: Technology at the Neural Interface 20, 290–300. 10.1111/ner.12541.

Yeo, B.T., Krienen, F.M., Sepulcre, J., Sabuncu, M.R., Lashkari, D., Hollinshead, M., Roffman, J.L., Smoller, J.W., Zollei, L., Polimeni, J.R., et al. (2011). The organization of the human cerebral cortex estimated by intrinsic functional connectivity. J Neurophysiol 106, 1125–1165. 10.1152/jn.00338.2011.

Zhang, Y., Brady, M., and Smith, S. (2001). Segmentation of brain MR images through a hidden Markov random field model and the expectation-maximization algorithm. IEEE Trans Med Imaging 20, 45–57. 10.1109/42.906424.

Zhang, Y., Lu, T., Dong, Y., Chen, Y., and Chen, J.D.Z. (2021). Auricular vagal nerve stimulation enhances gastrointestinal motility and improves interstitial cells of Cajal in rats treated with loperamide. Neurogastroenterol Motil, e14163. 10.1111/nmo.14163.

