## Supplementary Information for "Vagus nerve stimulation increases stomach-brain coupling via a vagal afferent pathway"

### Supplementary Information (SI)

Sophie Müller[1], Vanessa Teckentrup[1,6], Ignacio Rebollo[2], Manfred Hallschmid[3,4,5], Nils B. Kroemer*[1]

[1] University of Tübingen, Center for Mental Health, Department of Psychiatry and Psychotherapy, 72076 Tübingen, Germany

[2] German Institute of Human Nutrition (DIfE), Department of Decision Neuroscience and Nutrition (DNN), Potsdam-Rehbruecke, 14558 Nuthetal, Germany

[3] University of Tübingen, Department of Medical Psychology and Behavioral Neurobiology, 72076 Tübingen, Germany

[4] German Center for Diabetes Research (DZD), 85764 München-Neuherberg, Germany

[5] Institute for Diabetes Research and Metabolic Diseases of the Helmholtz Center Munich at the Eberhard Karls University Tübingen, 72076 Tübingen, Germany

[6] Trinity College Institute of Neuroscience, Trinity College Dublin, Dublin 2, Ireland

**Corresponding author***

Dr. Nils B. Kroemer,

Calwerstr. 14, 72076 Tübingen, Germany

Twitter: @cornu_copiae

### Supplementary Figures

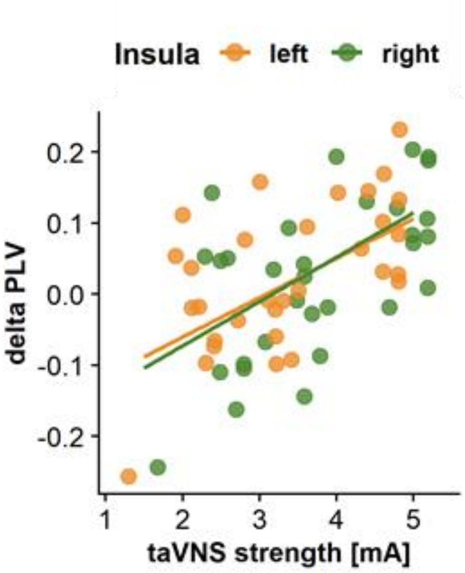


**Figure S1. Related to Figure 2.** We found strong positive correlations ($r$ = .58, $p$ < .001) between changes in PLV and taVNS strength in both insular cortices (clusters: right: $t_{max}$ = 4.52, $k$ = 102; left: $t_{max}$ = 4.10, $k$ = 190). This plot shows the extracted PLV changes in relation to the taVNS strength. Each point depicts one participant.

**Supplementary Tables**

**Table S1. Related to Figure 1. *t*-values during baseline.**

| | This study | | Rebollo (2018) | |
|---|---|---|---|---|
| **Region (Harvard Oxford Atlas, extended)** | **Mean *t*** | **Max *t*** | **Mean *t*** | **Max *t*** |
| FP r (Frontal Pole Right) | 0.918 | 5.104 | 1.125 | 5.074 |
| FP l (Frontal Pole Left) | 1.312 | 4.098 | 1.149 | 4.633 |
| IC r (Insular Cortex Right) | 1.705 | 4.809 | 1.421 | 4.473 |
| IC l (Insular Cortex Left) | 1.958 | 5.38 | 1.489 | 4.026 |
| SFG r (Superior Frontal Gyrus Right) | 1.133 | 6.487 | 1.204 | 4.895 |
| SFG l (Superior Frontal Gyrus Left) | 1.523 | 5.447 | 1.3 | 5.468 |
| MidFG r (Middle Frontal Gyrus Right) | 1.49 | 6.031 | 1.309 | 4.298 |
| MidFG l (Middle Frontal Gyrus Left) | 1.406 | 3.677 | 1.045 | 3.762 |
| IFG tri r (Inferior Frontal Gyrus, pars triangularis Right) | 1.895 | 4.458 | 1.306 | 3.864 |
| IFG tri l (Inferior Frontal Gyrus, pars triangularis Left) | 2.001 | 5.199 | 1.166 | 3.686 |
| IFG oper r (Inferior Frontal Gyrus, pars opercularis Right) | 2.209 | 5.933 | 1.411 | 3.946 |
| IFG oper l (Inferior Frontal Gyrus, pars opercularis Left) | 2.047 | 4.154 | 1.025 | 4.059 |
| PreCG r (Precentral Gyrus Right) | 2.362 | 5.819 | 1.677 | 4.525 |
| PreCG l (Precentral Gyrus Left) | 2.22 | 5.931 | 1.696 | 4.638 |
| TP r (Temporal Pole Right) | 1.14 | 5.176 | 1.045 | 4.669 |
| TP l (Temporal Pole Left) | 1.16 | 4.536 | 0.874 | 4.163 |
| aSTG r (Superior Temporal Gyrus, anterior division Right) | 1.743 | 3.746 | 1.636 | 4.014 |
| aSTG l (Superior Temporal Gyrus, anterior division Left) | 1.19 | 4.451 | 1.943 | 6.074 |
| pSTG r (Superior Temporal Gyrus, posterior division Right) | 1.316 | 2.998 | 1.77 | 3.933 |
| pSTG l (Superior Temporal Gyrus, posterior division Left) | 0.686 | 3.151 | 1.61 | 3.966 |
| aMTG r (Middle Temporal Gyrus, anterior division Right) | 1.14 | 2.554 | 1.284 | 4.153 |
| aMTG l (Middle Temporal Gyrus, anterior division Left) | 0.609 | 3.572 | 0.979 | 3.17 |
| pMTG r (Middle Temporal Gyrus, posterior division Right) | 0.807 | 3.3 | 1.305 | 4.153 |
| pMTG l (Middle Temporal Gyrus, posterior division Left) | 0.708 | 3.127 | 0.879 | 3.703 |
| toMTG r (Middle Temporal Gyrus, temporooccipital part Right) | 1.635 | 4.629 | 1.114 | 4.068 |
| toMTG l (Middle Temporal Gyrus, temporooccipital part Left) | 1.381 | 4.663 | 1.406 | 3.804 |
| aITG r (Inferior Temporal Gyrus, anterior division Right) | 2.094 | 4.42 | 1.436 | 3.631 |
| aITG l (Inferior Temporal Gyrus, anterior division Left) | 1.631 | 4.402 | 0.703 | 2.966 |

| Region (Harvard Oxford Atlas, extended) | This study Mean *t* | This study Max *t* | Rebollo (2018) Mean *t* | Rebollo (2018) Max *t* |
|---|---|---|---|---|
| pITG r (Inferior Temporal Gyrus, posterior division Right) | 1.223 | 3.902 | 0.925 | 3.648 |
| pITG l (Inferior Temporal Gyrus, posterior division Left) | 1.399 | 4.398 | 0.746 | 4.493 |
| toITG r (Inferior Temporal Gyrus, temporooccipital part Right) | 2.04 | 4.971 | 1.451 | 4.259 |
| toITG l (Inferior Temporal Gyrus, temporooccipital part Left) | 2.014 | 5.313 | 1.544 | 4.794 |
| PostCG r (Postcentral Gyrus Right) | 2.316 | 5.693 | 1.893 | 4.764 |
| PostCG l (Postcentral Gyrus Left) | 2.095 | 5.06 | 1.824 | 4.921 |
| SPL r (Superior Parietal Lobule Right) | 2.258 | 4.401 | 1.736 | 4.914 |
| SPL l (Superior Parietal Lobule Left) | 2.369 | 5.944 | 1.554 | 3.882 |
| aSMG r (Supramarginal Gyrus, anterior division Right) | 2.18 | 4.763 | 1.587 | 4.227 |
| aSMG l (Supramarginal Gyrus, anterior division Left) | 1.921 | 4.628 | 1.63 | 4.025 |
| pSMG r (Supramarginal Gyrus, posterior division Right) | 2.181 | 4.235 | 1.166 | 4.914 |
| pSMG l (Supramarginal Gyrus, posterior division Left) | 1.702 | 4.641 | 1.225 | 3.664 |
| AG r (Angular Gyrus Right) | 1.454 | 3.822 | 1.316 | 3.824 |
| AG l (Angular Gyrus Left) | 1.862 | 5.06 | 1.18 | 3.405 |
| sLOC r (Lateral Occipital Cortex, superior division Right) | 1.704 | 5.205 | 1.578 | 5.08 |
| sLOC l (Lateral Occipital Cortex, superior division Left) | 1.573 | 4.719 | 1.489 | 4.592 |
| iLOC r (Lateral Occipital Cortex, inferior division Right) | 2.204 | 4.664 | 1.713 | 4.479 |
| iLOC l (Lateral Occipital Cortex, inferior division Left) | 1.761 | 3.973 | 1.545 | 4.152 |
| ICC r (Intracalcarine Cortex Right) | 2.17 | 4.101 | 2.204 | 4.243 |
| ICC l (Intracalcarine Cortex Left) | 2.281 | 4.256 | 2.107 | 3.904 |
| MedFC (Frontal Medial Cortex) | 0.919 | 4.159 | 1.165 | 3.799 |
| SMA r (Juxtapositional Lobule Cortex- Ri | 2.154 | 5.043 | 2.132 | 4.753 |
| SMA L(Juxtapositional Lobule Cortex- Lef | 2.341 | 5.475 | 1.907 | 4.274 |
| SubCalC (Subcallosal Cortex) | 1.15 | 3.956 | 0.871 | 3.429 |
| PaCiG r (Paracingulate Gyrus Right) | 1.453 | 5.468 | 1.099 | 3.374 |
| PaCiG l (Paracingulate Gyrus Left) | 1.563 | 4.429 | 1.21 | 3.879 |
| AC (Cingulate Gyrus, anterior division) | 2.297 | 5.892 | 1.406 | 6.551 |
| PC (Cingulate Gyrus, posterior division) | 1.981 | 5.765 | 1.68 | 4.94 |
| Precuneous (Precuneous Cortex) | 2.153 | 5.792 | 1.801 | 4.859 |
| Cuneal r (Cuneal Cortex Right) | 1.605 | 3.916 | 2.407 | 4.863 |
| Cuneal l (Cuneal Cortex Left) | 1.815 | 3.422 | 2.34 | 4.464 |
| FOrb r (Frontal Orbital Cortex Right) | 1.032 | 4.099 | 1.328 | 4.195 |
| FOrb l (Frontal Orbital Cortex Left) | 1.28 | 5.074 | 1.392 | 4.543 |
| aPaHC r (Parahippocampal Gyrus, anterior division Right) | 1.649 | 6.599 | 1.331 | 3.826 |

| Region (Harvard Oxford Atlas, extended) | This study Mean *t* | This study Max *t* | Rebollo (2018) Mean *t* | Rebollo (2018) Max *t* |
|---|---|---|---|---|
| aPaHC l (Parahippocampal Gyrus, anterior division Left) | 1.56 | 5.067 | 1.067 | 3.625 |
| pPaHC r (Parahippocampal Gyrus, posterior division Right) | 1.912 | 3.826 | 1.104 | 3.76 |
| pPaHC l (Parahippocampal Gyrus, posterior division Left) | 1.465 | 3.966 | 0.936 | 3.632 |
| LG r (Lingual Gyrus Right) | 2.17 | 5.703 | 2.17 | 4.432 |
| LG l (Lingual Gyrus Left) | 1.802 | 4.555 | 2.131 | 4.948 |
| aTFusC r (Temporal Fusiform Cortex, anterior division Right) | 1.011 | 4.202 | 1.594 | 3.758 |
| aTFusC l (Temporal Fusiform Cortex, anterior division Left) | 1.236 | 3.343 | 0.957 | 3.988 |
| pTFusC r (Temporal Fusiform Cortex, posterior division Right) | 1.818 | 4.691 | 1.218 | 4.037 |
| pTFusC l (Temporal Fusiform Cortex, posterior division Left) | 1.561 | 3.929 | 1.063 | 3.673 |
| TOFusC r (Temporal Occipital Fusiform Cortex Right) | 2.555 | 6.842 | 1.748 | 4.243 |
| TOFusC l (Temporal Occipital Fusiform Cortex Left) | 2.16 | 4.111 | 1.827 | 4.617 |
| OFusG r (Occipital Fusiform Gyrus Right) | 2.563 | 5.394 | 1.965 | 5.206 |
| OFusG l (Occipital Fusiform Gyrus Left) | 2.571 | 5.185 | 2.026 | 4.136 |
| FO r (Frontal Operculum Cortex Right) | 1.951 | 4.22 | 1.417 | 3.763 |
| FO l (Frontal Operculum Cortex Left) | 1.969 | 4.209 | 1.145 | 3.67 |
| CO r (Central Opercular Cortex Right) | 2.628 | 5.149 | 1.988 | 4.763 |
| CO l (Central Opercular Cortex Left) | 2.181 | 5.215 | 1.731 | 4.112 |
| PO r (Parietal Operculum Cortex Right) | 2.193 | 4.32 | 1.938 | 4.209 |
| PO l (Parietal Operculum Cortex Left) | 2.544 | 5.018 | 2.095 | 4.084 |
| PP r (Planum Polare Right) | 1.992 | 4.328 | 1.59 | 4.006 |
| PP l (Planum Polare Left) | 1.97 | 5.203 | 1.067 | 3.08 |
| HG r (Heschl's Gyrus Right) | 2.412 | 4.686 | 2.111 | 5.174 |
| HG l (Heschl's Gyrus Left) | 2.165 | 5.051 | 2.131 | 5.169 |
| PT r (Planum Temporale Right) | 1.789 | 4.384 | 2.353 | 5.285 |
| PT l (Planum Temporale Left) | 1.653 | 4.264 | 1.998 | 4.481 |
| SCC r (Supracalcarine Cortex Right) | 2.093 | 3.735 | 2.246 | 3.966 |
| SCC l (Supracalcarine Cortex Left) | 2.299 | 4.107 | 2.208 | 3.839 |
| OP r (Occipital Pole Right) | 1.628 | 4.62 | 1.422 | 4.224 |
| OP l (Occipital Pole Left) | 1.638 | 4.018 | 1.221 | 4.074 |
| Thalamus r | 2.591 | 5.601 | 1.382 | 3.842 |
| Thalamus l | 2.199 | 5.274 | 1.461 | 4.673 |
| Caudate r | 0.887 | 3.136 | 1.838 | 4.9 |
| Caudate l | 1.365 | 4.386 | 1.511 | 4.326 |
| Putamen r | 1.413 | 3.802 | 1.21 | 4.184 |
| Putamen l | 1.453 | 4.286 | 1.401 | 4.825 |
| Pallidum r | 1.49 | 4.15 | 0.88 | 2.773 |
| Pallidum l | 1.842 | 5.182 | 0.918 | 4.057 |
| Hippocampus r | 1.586 | 4.293 | 1.191 | 3.727 |

| Region (Harvard Oxford Atlas, extended) | This study Mean *t* | This study Max *t* | Rebollo (2018) Mean *t* | Rebollo (2018) Max *t* |
|---|---|---|---|---|
| Hippocampus l | 1.649 | 4.618 | 0.838 | 3.465 |
| Amygdala r | 1.663 | 4.042 | 1.106 | 3.695 |
| Amygdala l | 1.618 | 3.405 | 0.969 | 3.625 |
| Accumbens r | 1.27 | 2.827 | 0.906 | 2.789 |
| Accumbens l | 0.741 | 2.172 | 1.556 | 2.936 |
| Brain-Stem | 1.578 | 6.262 | 1.039 | 4.514 |
| Cereb1 l (Cerebelum Crus1 Left) | 1.298 | 4.917 | 1.675 | 4.866 |
| Cereb1 r (Cerebelum Crus1 Right) | 1.53 | 4.959 | 1.417 | 4.82 |
| Cereb2 l (Cerebelum Crus2 Left) | 1.303 | 5.572 | 1.513 | 4.243 |
| Cereb2 r (Cerebelum Crus2 Right) | 1.778 | 4.808 | 1.13 | 4.82 |
| Cereb3 l (Cerebelum 3 Left) | 1.503 | 3.538 | 1.265 | 3.04 |
| Cereb3 r (Cerebelum 3 Right) | 1.658 | 4.011 | 1.338 | 3.592 |
| Cereb45 l (Cerebelum 4 5 Left) | 1.626 | 4.355 | 1.437 | 4.72 |
| Cereb45 r (Cerebelum 4 5 Right) | 1.683 | 3.88 | 1.485 | 4.003 |
| Cereb6 l (Cerebelum 6 Left) | 1.802 | 4.683 | 1.669 | 4.59 |
| Cereb6 r (Cerebelum 6 Right) | 2.004 | 6.336 | 1.831 | 4.483 |
| Cereb7 l (Cerebelum 7b Left) | 1.881 | 4.871 | 0.869 | 4.387 |
| Cereb7 r (Cerebelum 7b Right) | 2.063 | 5.825 | 0.627 | 3.936 |
| Cereb8 l (Cerebelum 8 Left) | 1.728 | 5.49 | 0.601 | 3.591 |
| Cereb8 r (Cerebelum 8 Right) | 1.85 | 5.522 | 0.592 | 4.503 |
| Cereb9 l (Cerebelum 9 Left) | 1.61 | 4.825 | 0.793 | 3.397 |
| Cereb9 r (Cerebelum 9 Right) | 1.787 | 5.12 | 0.786 | 3.752 |
| Cereb10 l (Cerebelum 10 Left) | 1.482 | 3.932 | 1.676 | 4.387 |
| Cereb10 r (Cerebelum 10 Right) | 2.252 | 5.251 | 1.468 | 2.874 |
| Ver12 (Vermis 1 2) | 0.722 | 2.147 | 1.42 | 2.733 |
| Ver3 (Vermis 3) | 2.339 | 5.212 | 1.026 | 2.733 |
| Ver45 (Vermis 4 5) | 2.071 | 5.204 | 2.062 | 4.648 |
| Ver6 (Vermis 6) | 2.372 | 4.831 | 1.875 | 3.455 |
| Ver7 (Vermis 7) | 2.233 | 4.619 | 1.984 | 3.993 |
| Ver8 (Vermis 8) | 2.821 | 5.331 | 1.869 | 3.641 |
| Ver9 (Vermis 9) | 1.538 | 3.272 | 1.856 | 3.621 |
| Ver10 (Vermis 10) | 1.484 | 3.478 | 0.749 | 2.834 |
| Extended amygdala r | 1.492 | 2.814 | 1.772 | 3.277 |
| Extended amygdala l | 0.805 | 2.524 | 0.926 | 2.469 |
| Substantia nigra pars compacta r | 2.074 | 4.026 | 0.649 | 3.198 |
| Substantia nigra pars compacta l | 1.934 | 3.382 | 0.571 | 2.23 |
| Nucleus ruber r | 1.317 | 3.745 | 1.493 | 2.454 |
| Nucleus ruber l | 1.649 | 3.038 | 0.783 | 2.419 |
| Substantia nigra pars reticularis r | 2.261 | 4.785 | 0.915 | 3.198 |
| Substantia nigra pars reticularis l | 1.877 | 2.969 | 0.716 | 2.568 |
| Parabrachial pigmented nucleus r | 1.683 | 3.257 | 0.823 | 3.198 |
| Parabrachial pigmented nucleus l | 2.03 | 3.352 | 0.481 | 1.851 |
| Ventral tegmental area r | 2.297 | 3.138 | 0.942 | 1.755 |
| Ventral tegmental area l | 1.499 | 2.507 | 0.95 | 1.851 |
| Ventral pallidum r | 0.994 | 1.736 | 0.549 | 1.528 |
| Ventral pallidum l | 0.65 | 2.519 | 1.504 | 2.555 |
| Hypothalamus r | 1.349 | 3.124 | 0.329 | 3.569 |

| Region (Harvard Oxford Atlas, extended) | This study Mean *t* | Max *t* | Rebollo (2018) Mean *t* | Max *t* |
|---|---|---|---|---|
| Hypothalamus l | 1.602 | 3.754 | 1.033 | 2.383 |
| Mammilary body r | 1.062 | 2.335 | 2.594 | 3.368 |
| Mammilary body l | 1.949 | 2.872 | 1.509 | 2.752 |
| Subthalamic nucleus r | 1.89 | 2.668 | 0.624 | 1.903 |
| Subthalamic nucleus l | 2.253 | 3.902 | 0.305 | 2.383 |
| NTS new r | 1.252 | 2.842 | 0.136 | 1.905 |
| NTS new l | 1.191 | 3.006 | 0.081 | 2.704 |

**Table S2. Related to Figure 2. Effect sizes per region.**

| Region (Harvard Oxford Atlas, extended) | Baseline Mean $d_z$ | Baseline Max $d_z$ | Baseline Std $d_z$ | taVNS > Sham Mean $d_z$ | taVNS > Sham Max $d_z$ | taVNS > Sham Std $d_z$ |
|---|---|---|---|---|---|---|
| FP r (Frontal Pole Right) | 0.17 | 0.917 | 0.188 | 0.014 | 0.535 | 0.154 |
| FP l (Frontal Pole Left) | 0.247 | 0.736 | 0.161 | 0.024 | 0.765 | 0.158 |
| IC r (Insular Cortex Right) | 0.306 | 0.864 | 0.187 | 0.112 | 0.59 | 0.16 |
| IC l (Insular Cortex Left) | 0.352 | 0.966 | 0.186 | 0.119 | 0.609 | 0.163 |
| SFG r (Superior Frontal Gyrus Right) | 0.204 | 1.165 | 0.239 | 0.045 | 0.66 | 0.175 |
| SFG l (Superior Frontal Gyrus Left) | 0.273 | 0.978 | 0.171 | -0.02 | 0.471 | 0.148 |
| MidFG r (Middle Frontal Gyrus Right) | 0.268 | 1.083 | 0.201 | 0.101 | 0.893 | 0.202 |
| MidFG l (Middle Frontal Gyrus Left) | 0.252 | 0.66 | 0.142 | 0.041 | 0.511 | 0.168 |
| IFG tri r (Inferior Frontal Gyrus, pars triangularis Right) | 0.34 | 0.801 | 0.146 | 0.033 | 0.62 | 0.175 |
| IFG tri l (Inferior Frontal Gyrus, pars triangularis Left) | 0.36 | 0.934 | 0.148 | -0.036 | 0.422 | 0.139 |
| IFG oper r (Inferior Frontal Gyrus, pars opercularis Right) | 0.397 | 1.066 | 0.2 | 0.087 | 0.573 | 0.175 |
| IFG oper l (Inferior Frontal Gyrus, pars opercularis Left) | 0.368 | 0.746 | 0.131 | -0.106 | 0.389 | 0.152 |
| PreCG r (Precentral Gyrus Right) | 0.424 | 1.045 | 0.154 | -0.023 | 0.612 | 0.16 |
| PreCG l (Precentral Gyrus Left) | 0.401 | 1.065 | 0.155 | -0.018 | 0.58 | 0.165 |
| TP r (Temporal Pole Right) | 0.211 | 0.93 | 0.169 | 0.009 | 0.751 | 0.174 |
| TP l (Temporal Pole Left) | 0.216 | 0.815 | 0.175 | -0.064 | 0.606 | 0.187 |
| aSTG r (Superior Temporal Gyrus, anterior division Right) | 0.313 | 0.673 | 0.128 | 0.067 | 0.462 | 0.114 |
| aSTG l (Superior Temporal Gyrus, anterior division Left) | 0.214 | 0.799 | 0.135 | -0.114 | 0.297 | 0.131 |
| pSTG r (Superior Temporal Gyrus, posterior division Right) | 0.236 | 0.538 | 0.133 | -0.08 | 0.348 | 0.157 |
| pSTG l (Superior Temporal Gyrus, posterior division Left) | 0.127 | 0.566 | 0.141 | -0.155 | 0.2 | 0.137 |
| aMTG r (Middle Temporal Gyrus, anterior division Right) | 0.206 | 0.459 | 0.094 | 0.098 | 0.411 | 0.119 |
| aMTG l (Middle Temporal Gyrus, anterior division Left) | 0.113 | 0.642 | 0.102 | -0.015 | 0.439 | 0.15 |
| pMTG r (Middle Temporal Gyrus, posterior division Right) | 0.146 | 0.593 | 0.185 | 0.087 | 0.68 | 0.169 |
| pMTG l (Middle Temporal Gyrus, posterior division Left) | 0.139 | 0.562 | 0.143 | -0.04 | 0.527 | 0.186 |
| toMTG r (Middle Temporal Gyrus, temporooccipital part Right) | 0.302 | 0.831 | 0.156 | 0.165 | 0.612 | 0.176 |
| toMTG l (Middle Temporal Gyrus, temporooccipital part Left) | 0.274 | 0.838 | 0.213 | 0.012 | 0.467 | 0.162 |
| aITG r (Inferior Temporal Gyrus, anterior division Right) | 0.377 | 0.794 | 0.204 | 0.076 | 0.462 | 0.151 |

| Region (Harvard Oxford Atlas, extended) | Baseline | | | taVNS > Sham | | |
|---|---|---|---|---|---|---|
| | Mean $d_z$ | Max $d_z$ | Std $d_z$ | Mean $d_z$ | Max $d_z$ | Std $d_z$ |
| aITG l (Inferior Temporal Gyrus, anterior division Left) | 0.309 | 0.791 | 0.196 | -0.024 | 0.565 | 0.149 |
| pITG r (Inferior Temporal Gyrus, posterior division Right) | 0.226 | 0.701 | 0.131 | 0.127 | 0.631 | 0.153 |
| pITG l (Inferior Temporal Gyrus, posterior division Left) | 0.256 | 0.79 | 0.149 | 0.08 | 0.578 | 0.14 |
| toITG r (Inferior Temporal Gyrus, temporooccipital part Right) | 0.371 | 0.893 | 0.221 | 0.096 | 0.608 | 0.171 |
| toITG l (Inferior Temporal Gyrus, temporooccipital part Left) | 0.373 | 0.954 | 0.183 | 0.098 | 0.485 | 0.143 |
| PostCG r (Postcentral Gyrus Right) | 0.416 | 1.023 | 0.142 | 0.001 | 0.494 | 0.164 |
| PostCG l (Postcentral Gyrus Left) | 0.379 | 0.909 | 0.146 | -0.053 | 0.484 | 0.165 |
| SPL r (Superior Parietal Lobule Right) | 0.405 | 0.791 | 0.152 | -0.014 | 0.487 | 0.157 |
| SPL l (Superior Parietal Lobule Left) | 0.426 | 1.068 | 0.174 | -0.039 | 0.412 | 0.162 |
| aSMG r (Supramarginal Gyrus, anterior division Right) | 0.392 | 0.855 | 0.128 | 0.118 | 0.496 | 0.126 |
| aSMG l (Supramarginal Gyrus, anterior division Left) | 0.354 | 0.831 | 0.16 | 0.145 | 0.55 | 0.139 |
| pSMG r (Supramarginal Gyrus, posterior division Right) | 0.397 | 0.761 | 0.129 | 0.134 | 0.631 | 0.163 |
| pSMG l (Supramarginal Gyrus, posterior division Left) | 0.322 | 0.834 | 0.169 | 0.015 | 0.444 | 0.151 |
| AG r (Angular Gyrus Right) | 0.269 | 0.686 | 0.147 | 0.15 | 0.55 | 0.133 |
| AG l (Angular Gyrus Left) | 0.349 | 0.909 | 0.15 | 0.096 | 0.481 | 0.168 |
| sLOC r (Lateral Occipital Cortex, superior division Right) | 0.328 | 0.935 | 0.188 | 0.128 | 0.763 | 0.198 |
| sLOC l (Lateral Occipital Cortex, superior division Left) | 0.315 | 0.847 | 0.164 | 0.101 | 0.508 | 0.145 |
| iLOC r (Lateral Occipital Cortex, inferior division Right) | 0.434 | 0.838 | 0.114 | -0.056 | 0.363 | 0.12 |
| iLOC l (Lateral Occipital Cortex, inferior division Left) | 0.373 | 0.714 | 0.139 | -0.084 | 0.452 | 0.147 |
| ICC r (Intracalcarine Cortex Right) | 0.39 | 0.736 | 0.136 | 0.12 | 0.355 | 0.094 |
| ICC l (Intracalcarine Cortex Left) | 0.41 | 0.764 | 0.118 | 0.088 | 0.406 | 0.091 |
| MedFC (Frontal Medial Cortex) | 0.165 | 0.747 | 0.167 | 0.077 | 0.539 | 0.148 |
| SMA r (Juxtapositional Lobule Cortex- Ri | 0.387 | 0.906 | 0.16 | 0.044 | 0.413 | 0.131 |
| SMA L(Juxtapositional Lobule Cortex- Lef | 0.421 | 0.983 | 0.142 | 0.014 | 0.459 | 0.134 |
| SubCalC (Subcallosal Cortex) | 0.206 | 0.71 | 0.17 | 0.169 | 0.676 | 0.175 |
| PaCiG r (Paracingulate Gyrus Right) | 0.261 | 0.982 | 0.212 | 0.163 | 0.654 | 0.144 |

| Region (Harvard Oxford Atlas, extended) | Baseline Mean $d_z$ | Baseline Max $d_z$ | Baseline Std $d_z$ | taVNS > Sham Mean $d_z$ | taVNS > Sham Max $d_z$ | taVNS > Sham Std $d_z$ |
|---|---|---|---|---|---|---|
| PaCiG l (Paracingulate Gyrus Left) | 0.281 | 0.796 | 0.166 | 0.127 | 0.75 | 0.153 |
| AC (Cingulate Gyrus, anterior division) | 0.412 | 1.058 | 0.19 | 0.141 | 0.797 | 0.174 |
| PC (Cingulate Gyrus, posterior division) | 0.356 | 1.035 | 0.158 | 0.228 | 0.699 | 0.149 |
| Precuneous (Precuneous Cortex) | 0.387 | 1.04 | 0.126 | 0.169 | 0.762 | 0.17 |
| Cuneal r (Cuneal Cortex Right) | 0.288 | 0.703 | 0.119 | 0.069 | 0.392 | 0.111 |
| Cuneal l (Cuneal Cortex Left) | 0.326 | 0.615 | 0.076 | 0.045 | 0.385 | 0.117 |
| FOrb r (Frontal Orbital Cortex Right) | 0.186 | 0.736 | 0.168 | 0.078 | 0.709 | 0.189 |
| FOrb l (Frontal Orbital Cortex Left) | 0.231 | 0.911 | 0.167 | 0.116 | 0.604 | 0.173 |
| aPaHC r (Parahippocampal Gyrus, anterior division Right) | 0.298 | 1.185 | 0.22 | 0.045 | 0.518 | 0.159 |
| aPaHC l (Parahippocampal Gyrus, anterior division Left) | 0.281 | 0.91 | 0.152 | 0.02 | 0.398 | 0.148 |
| pPaHC r (Parahippocampal Gyrus, posterior division Right) | 0.343 | 0.687 | 0.139 | 0.06 | 0.459 | 0.178 |
| pPaHC l (Parahippocampal Gyrus, posterior division Left) | 0.263 | 0.712 | 0.163 | 0.042 | 0.395 | 0.174 |
| LG r (Lingual Gyrus Right) | 0.39 | 1.024 | 0.158 | 0.064 | 0.469 | 0.14 |
| LG l (Lingual Gyrus Left) | 0.324 | 0.818 | 0.144 | 0.095 | 0.608 | 0.15 |
| aTFusC r (Temporal Fusiform Cortex, anterior division Right) | 0.185 | 0.755 | 0.221 | 0.076 | 0.411 | 0.114 |
| aTFusC l (Temporal Fusiform Cortex, anterior division Left) | 0.23 | 0.6 | 0.155 | -0.065 | 0.294 | 0.14 |
| pTFusC r (Temporal Fusiform Cortex, posterior division Right) | 0.341 | 0.843 | 0.146 | 0.147 | 0.553 | 0.151 |
| pTFusC l (Temporal Fusiform Cortex, posterior division Left) | 0.285 | 0.706 | 0.138 | -0.004 | 0.554 | 0.181 |
| TOFusC r (Temporal Occipital Fusiform Cortex Right) | 0.459 | 1.229 | 0.155 | 0.089 | 0.548 | 0.139 |
| TOFusC l (Temporal Occipital Fusiform Cortex Left) | 0.388 | 0.738 | 0.144 | 0.14 | 0.528 | 0.146 |
| OFusG r (Occipital Fusiform Gyrus Right) | 0.46 | 0.969 | 0.158 | 0.027 | 0.405 | 0.113 |
| OFusG l (Occipital Fusiform Gyrus Left) | 0.462 | 0.931 | 0.107 | 0.1 | 0.387 | 0.117 |
| FO r (Frontal Operculum Cortex Right) | 0.35 | 0.758 | 0.159 | 0.118 | 0.523 | 0.149 |
| FO l (Frontal Operculum Cortex Left) | 0.354 | 0.756 | 0.136 | 0.162 | 0.594 | 0.193 |
| CO r (Central Opercular Cortex Right) | 0.472 | 0.925 | 0.141 | 0.176 | 0.551 | 0.136 |
| CO l (Central Opercular Cortex Left) | 0.392 | 0.937 | 0.171 | 0.113 | 0.549 | 0.131 |

| Region (Harvard Oxford Atlas, extended) | Baseline Mean $d_z$ | Baseline Max $d_z$ | Baseline Std $d_z$ | taVNS > Sham Mean $d_z$ | taVNS > Sham Max $d_z$ | taVNS > Sham Std $d_z$ |
|---|---|---|---|---|---|---|
| PO r (Parietal Operculum Cortex Right) | 0.394 | 0.776 | 0.15 | 0.048 | 0.335 | 0.119 |
| PO l (Parietal Operculum Cortex Left) | 0.457 | 0.901 | 0.139 | -0.009 | 0.519 | 0.141 |
| PP r (Planum Polare Right) | 0.358 | 0.777 | 0.18 | 0.022 | 0.63 | 0.178 |
| PP l (Planum Polare Left) | 0.354 | 0.935 | 0.167 | -0.028 | 0.633 | 0.211 |
| HG r (Heschl's Gyrus Right) | 0.433 | 0.842 | 0.141 | 0.171 | 0.511 | 0.133 |
| HG l (Heschl's Gyrus Left) | 0.389 | 0.907 | 0.149 | 0.076 | 0.376 | 0.118 |
| PT r (Planum Temporale Right) | 0.321 | 0.787 | 0.142 | 0.023 | 0.477 | 0.138 |
| PT l (Planum Temporale Left) | 0.297 | 0.766 | 0.174 | -0.099 | 0.36 | 0.175 |
| SCC r (Supracalcarine Cortex Right) | 0.376 | 0.671 | 0.138 | 0.131 | 0.344 | 0.089 |
| SCC l (Supracalcarine Cortex Left) | 0.413 | 0.738 | 0.122 | 0.091 | 0.365 | 0.115 |
| OP r (Occipital Pole Right) | 0.352 | 0.83 | 0.136 | 0.065 | 0.367 | 0.111 |
| OP l (Occipital Pole Left) | 0.373 | 0.722 | 0.131 | 0.078 | 0.443 | 0.14 |
| Thalamus r | 0.465 | 1.006 | 0.163 | 0.038 | 0.614 | 0.141 |
| Thalamus l | 0.395 | 0.947 | 0.17 | -0.008 | 0.444 | 0.153 |
| Caudate r | 0.159 | 0.563 | 0.146 | -0.07 | 0.327 | 0.12 |
| Caudate l | 0.245 | 0.788 | 0.192 | -0.052 | 0.234 | 0.124 |
| Putamen r | 0.254 | 0.683 | 0.148 | -0.165 | 0.308 | 0.173 |
| Putamen l | 0.261 | 0.77 | 0.155 | -0.158 | 0.229 | 0.152 |
| Pallidum r | 0.268 | 0.745 | 0.177 | 0.095 | 0.49 | 0.158 |
| Pallidum l | 0.331 | 0.931 | 0.186 | 0.096 | 0.542 | 0.182 |
| Hippocampus r | 0.285 | 0.771 | 0.161 | -0.01 | 0.445 | 0.154 |
| Hippocampus l | 0.296 | 0.829 | 0.164 | 0.026 | 0.517 | 0.167 |
| Amygdala r | 0.299 | 0.726 | 0.136 | -0.112 | 0.294 | 0.138 |
| Amygdala l | 0.291 | 0.612 | 0.121 | -0.032 | 0.474 | 0.151 |
| Accumbens r | 0.228 | 0.508 | 0.123 | 0.076 | 0.345 | 0.136 |
| Accumbens l | 0.133 | 0.39 | 0.149 | 0.107 | 0.362 | 0.12 |
| Brain-Stem | 0.289 | 1.125 | 0.184 | 0.012 | 0.549 | 0.173 |
| Cereb1 l (Cerebelum Crus1 Left) | 0.252 | 0.883 | 0.156 | 0.121 | 0.564 | 0.143 |
| Cereb1 r (Cerebelum Crus1 Right) | 0.297 | 0.891 | 0.155 | 0.095 | 0.484 | 0.128 |
| Cereb2 l (Cerebelum Crus2 Left) | 0.256 | 1.001 | 0.179 | 0.113 | 0.457 | 0.113 |
| Cereb2 r (Cerebelum Crus2 Right) | 0.364 | 0.864 | 0.144 | 0.102 | 0.569 | 0.125 |
| Cereb3 l (Cerebelum 3 Left) | 0.27 | 0.635 | 0.138 | 0.26 | 0.607 | 0.155 |
| Cereb3 r (Cerebelum 3 Right) | 0.298 | 0.72 | 0.152 | 0.095 | 0.453 | 0.158 |
| Cereb45 l (Cerebelum 4 5 Left) | 0.294 | 0.782 | 0.151 | 0.117 | 0.579 | 0.164 |
| Cereb45 r (Cerebelum 4 5 Right) | 0.306 | 0.697 | 0.135 | 0.12 | 0.65 | 0.164 |
| Cereb6 l (Cerebelum 6 Left) | 0.331 | 0.841 | 0.174 | 0.153 | 0.533 | 0.154 |
| Cereb6 r (Cerebelum 6 Right) | 0.374 | 1.138 | 0.164 | 0.121 | 0.86 | 0.171 |
| Cereb7 l (Cerebelum 7b Left) | 0.376 | 0.875 | 0.173 | 0.088 | 0.444 | 0.133 |
| Cereb7 r (Cerebelum 7b Right) | 0.412 | 1.046 | 0.207 | 0.075 | 0.405 | 0.111 |
| Cereb8 l (Cerebelum 8 Left) | 0.321 | 0.986 | 0.177 | 0.059 | 0.476 | 0.143 |
| Cereb8 r (Cerebelum 8 Right) | 0.343 | 0.992 | 0.189 | 0.108 | 0.622 | 0.139 |
| Cereb9 l (Cerebelum 9 Left) | 0.289 | 0.867 | 0.192 | 0.049 | 0.697 | 0.167 |

| Region (Harvard Oxford Atlas, extended) | Baseline | | | taVNS > Sham | | |
|---|---|---|---|---|---|---|
| | Mean $d_z$ | Max $d_z$ | Std $d_z$ | Mean $d_z$ | Max $d_z$ | Std $d_z$ |
| Cereb9 r (Cerebelum 9 Right) | 0.321 | 0.92 | 0.189 | 0.044 | 0.454 | 0.161 |
| Cereb10 l (Cerebelum 10 Left) | 0.287 | 0.706 | 0.143 | 0.054 | 0.443 | 0.133 |
| Cereb10 r (Cerebelum 10 Right) | 0.497 | 0.943 | 0.186 | 0.136 | 0.441 | 0.106 |
| Ver12 (Vermis 1 2) | 0.13 | 0.386 | 0.153 | -0.04 | 0.329 | 0.165 |
| Ver3 (Vermis 3) | 0.42 | 0.936 | 0.221 | 0.202 | 0.556 | 0.155 |
| Ver45 (Vermis 4 5) | 0.372 | 0.935 | 0.168 | 0.108 | 0.603 | 0.186 |
| Ver6 (Vermis 6) | 0.426 | 0.868 | 0.154 | 0.09 | 0.529 | 0.196 |
| Ver7 (Vermis 7) | 0.401 | 0.83 | 0.136 | 0.082 | 0.4 | 0.122 |
| Ver8 (Vermis 8) | 0.507 | 0.957 | 0.197 | 0.018 | 0.466 | 0.139 |
| Ver9 (Vermis 9) | 0.276 | 0.588 | 0.12 | 0.17 | 0.623 | 0.125 |
| Ver10 (Vermis 10) | 0.267 | 0.625 | 0.144 | 0.073 | 0.495 | 0.195 |
| Extended amygdala r | 0.268 | 0.505 | 0.103 | 0.027 | 0.177 | 0.097 |
| Extended amygdala l | 0.145 | 0.453 | 0.148 | -0.038 | 0.163 | 0.15 |
| Substantia nigra pars compacta r | 0.372 | 0.723 | 0.169 | 0.013 | 0.233 | 0.157 |
| Substantia nigra pars compacta l | 0.347 | 0.608 | 0.121 | 0.139 | 0.419 | 0.137 |
| Nucleus ruber r | 0.237 | 0.673 | 0.172 | 0.233 | 0.531 | 0.124 |
| Nucleus ruber l | 0.296 | 0.546 | 0.11 | 0.08 | 0.547 | 0.157 |
| Substantia nigra pars reticularis r | 0.406 | 0.859 | 0.167 | 0.07 | 0.423 | 0.168 |
| Substantia nigra pars reticularis l | 0.337 | 0.533 | 0.144 | 0.131 | 0.503 | 0.129 |
| Parabrachial pigmented nucleus r | 0.302 | 0.585 | 0.192 | 0.142 | 0.405 | 0.139 |
| Parabrachial pigmented nucleus l | 0.365 | 0.602 | 0.138 | 0.038 | 0.325 | 0.137 |
| Ventral tegmental area r | 0.413 | 0.564 | 0.112 | 0.084 | 0.237 | 0.112 |
| Ventral tegmental area l | 0.269 | 0.45 | 0.116 | 0.343 | 0.611 | 0.223 |
| Ventral pallidum r | 0.179 | 0.312 | 0.063 | 0.01 | 0.119 | 0.075 |
| Ventral pallidum l | 0.117 | 0.453 | 0.167 | -0.162 | 0.134 | 0.134 |
| Hypothalamus r | 0.242 | 0.561 | 0.156 | 0.032 | 0.483 | 0.192 |
| Hypothalamus l | 0.288 | 0.674 | 0.192 | 0.053 | 0.36 | 0.168 |
| Mammilary body r | 0.191 | 0.419 | 0.133 | 0.045 | 0.179 | 0.114 |
| Mammilary body l | 0.35 | 0.516 | 0.105 | -0.059 | 0.159 | 0.123 |
| Subthalamic nucleus r | 0.339 | 0.479 | 0.089 | 0.224 | 0.551 | 0.149 |
| Subthalamic nucleus l | 0.405 | 0.701 | 0.174 | 0.194 | 0.468 | 0.175 |
| NTS new r | 0.225 | 0.51 | 0.134 | 0.055 | 0.75 | 0.258 |
| NTS new l | 0.214 | 0.54 | 0.129 | 0.221 | 0.778 | 0.208 |

**Table S3. Related to Figure 4. Correlation between changes in PLV and changes in hunger.**

| Region (Harvard Oxford Atlas, extended) | Mean *r* | Max *r* | Std *r* |
|---|---|---|---|
| FP r (Frontal Pole Right) | 0.117 | 0.764 | 0.197 |
| FP l (Frontal Pole Left) | 0.159 | 0.686 | 0.204 |
| IC r (Insular Cortex Right) | -0.11 | 0.359 | 0.163 |
| IC l (Insular Cortex Left) | -0.085 | 0.351 | 0.16 |
| SFG r (Superior Frontal Gyrus Right) | 0.064 | 0.546 | 0.158 |
| SFG l (Superior Frontal Gyrus Left) | 0.14 | 0.622 | 0.203 |
| MidFG r (Middle Frontal Gyrus Right) | 0.051 | 0.512 | 0.193 |
| MidFG l (Middle Frontal Gyrus Left) | 0.174 | 0.617 | 0.178 |
| IFG tri r (Inferior Frontal Gyrus, pars triangularis Right) | 0.025 | 0.497 | 0.151 |
| IFG tri l (Inferior Frontal Gyrus, pars triangularis Left) | 0.141 | 0.518 | 0.134 |
| IFG oper r (Inferior Frontal Gyrus, pars opercularis Right) | -0.102 | 0.339 | 0.169 |
| IFG oper l (Inferior Frontal Gyrus, pars opercularis Left) | -0.009 | 0.409 | 0.18 |
| PreCG r (Precentral Gyrus Right) | 0.069 | 0.604 | 0.185 |
| PreCG l (Precentral Gyrus Left) | 0.034 | 0.547 | 0.164 |
| TP r (Temporal Pole Right) | 0.126 | 0.536 | 0.16 |
| TP l (Temporal Pole Left) | 0.11 | 0.566 | 0.162 |
| aSTG r (Superior Temporal Gyrus, anterior division Right) | 0.014 | 0.424 | 0.157 |
| aSTG l (Superior Temporal Gyrus, anterior division Left) | -0.037 | 0.325 | 0.167 |
| pSTG r (Superior Temporal Gyrus, posterior division Right) | 0.017 | 0.417 | 0.159 |
| pSTG l (Superior Temporal Gyrus, posterior division Left) | -0.024 | 0.361 | 0.164 |
| aMTG r (Middle Temporal Gyrus, anterior division Right) | 0.116 | 0.503 | 0.129 |
| aMTG l (Middle Temporal Gyrus, anterior division Left) | 0.121 | 0.462 | 0.148 |
| pMTG r (Middle Temporal Gyrus, posterior division Right) | 0.243 | 0.616 | 0.138 |
| pMTG l (Middle Temporal Gyrus, posterior division Left) | 0.172 | 0.558 | 0.162 |
| toMTG r (Middle Temporal Gyrus, temporooccipital part Right) | 0.043 | 0.514 | 0.214 |
| toMTG l (Middle Temporal Gyrus, temporooccipital part Left) | 0.045 | 0.528 | 0.173 |
| aITG r (Inferior Temporal Gyrus, anterior division Right) | -0.003 | 0.385 | 0.169 |
| aITG l (Inferior Temporal Gyrus, anterior division Left) | 0.087 | 0.434 | 0.145 |
| pITG r (Inferior Temporal Gyrus, posterior division Right) | 0.105 | 0.541 | 0.176 |
| pITG l (Inferior Temporal Gyrus, posterior division Left) | 0.062 | 0.529 | 0.164 |

| Region (Harvard Oxford Atlas, extended) | Mean *r* | Max *r* | Std *r* |
|---|---|---|---|
| toITG r (Inferior Temporal Gyrus, temporooccipital part Right) | 0.301 | 0.622 | 0.148 |
| toITG l (Inferior Temporal Gyrus, temporooccipital part Left) | 0.345 | 0.627 | 0.147 |
| PostCG r (Postcentral Gyrus Right) | 0.098 | 0.611 | 0.17 |
| PostCG l (Postcentral Gyrus Left) | 0.094 | 0.491 | 0.157 |
| SPL r (Superior Parietal Lobule Right) | 0.045 | 0.519 | 0.19 |
| SPL l (Superior Parietal Lobule Left) | 0.121 | 0.568 | 0.217 |
| aSMG r (Supramarginal Gyrus, anterior division Right) | -0.132 | 0.313 | 0.155 |
| aSMG l (Supramarginal Gyrus, anterior division Left) | 0.058 | 0.461 | 0.199 |
| pSMG r (Supramarginal Gyrus, posterior division Right) | 0.027 | 0.51 | 0.173 |
| pSMG l (Supramarginal Gyrus, posterior division Left) | 0.026 | 0.384 | 0.16 |
| AG r (Angular Gyrus Right) | 0.115 | 0.555 | 0.203 |
| AG l (Angular Gyrus Left) | 0.222 | 0.602 | 0.158 |
| sLOC r (Lateral Occipital Cortex, superior division Right) | 0.214 | 0.634 | 0.177 |
| sLOC l (Lateral Occipital Cortex, superior division Left) | 0.219 | 0.638 | 0.174 |
| iLOC r (Lateral Occipital Cortex, inferior division Right) | 0.25 | 0.561 | 0.173 |
| iLOC l (Lateral Occipital Cortex, inferior division Left) | 0.243 | 0.627 | 0.155 |
| ICC r (Intracalcarine Cortex Right) | 0.045 | 0.356 | 0.117 |
| ICC l (Intracalcarine Cortex Left) | 0.11 | 0.486 | 0.145 |
| MedFC (Frontal Medial Cortex) | 0.116 | 0.523 | 0.156 |
| SMA r (Juxtapositional Lobule Cortex- Ri | 0.039 | 0.458 | 0.136 |
| SMA L(Juxtapositional Lobule Cortex- Lef | 0.06 | 0.39 | 0.134 |
| SubCalC (Subcallosal Cortex) | 0.071 | 0.5 | 0.149 |
| PaCiG r (Paracingulate Gyrus Right) | -0.089 | 0.483 | 0.192 |
| PaCiG l (Paracingulate Gyrus Left) | 0.024 | 0.56 | 0.177 |
| AC (Cingulate Gyrus, anterior division) | -0.139 | 0.389 | 0.141 |
| PC (Cingulate Gyrus, posterior division) | 0.119 | 0.608 | 0.216 |
| Precuneous (Precuneous Cortex) | 0.138 | 0.622 | 0.179 |
| Cuneal r (Cuneal Cortex Right) | -0.082 | 0.356 | 0.096 |
| Cuneal l (Cuneal Cortex Left) | -0.11 | 0.184 | 0.129 |
| FOrb r (Frontal Orbital Cortex Right) | 0.077 | 0.48 | 0.171 |
| FOrb l (Frontal Orbital Cortex Left) | 0.009 | 0.417 | 0.158 |
| aPaHC r (Parahippocampal Gyrus, anterior division Right) | 0.101 | 0.536 | 0.178 |
| aPaHC l (Parahippocampal Gyrus, anterior division Left) | 0.041 | 0.445 | 0.161 |
| pPaHC r (Parahippocampal Gyrus, posterior division Right) | 0.034 | 0.445 | 0.172 |
| pPaHC l (Parahippocampal Gyrus, posterior division Left) | 0.079 | 0.424 | 0.138 |
| LG r (Lingual Gyrus Right) | 0.056 | 0.473 | 0.142 |
| LG l (Lingual Gyrus Left) | 0.07 | 0.461 | 0.137 |

| **Region (Harvard Oxford Atlas, extended)** | **Mean *r*** | **Max *r*** | **Std *r*** |
|---|---|---|---|
| aTFusC r (Temporal Fusiform Cortex, anterior division Right) | 0.135 | 0.509 | 0.136 |
| aTFusC l (Temporal Fusiform Cortex, anterior division Left) | -0.024 | 0.299 | 0.131 |
| pTFusC r (Temporal Fusiform Cortex, posterior division Right) | 0.105 | 0.52 | 0.157 |
| pTFusC l (Temporal Fusiform Cortex, posterior division Left) | 0.111 | 0.456 | 0.148 |
| TOFusC r (Temporal Occipital Fusiform Cortex Right) | 0.269 | 0.645 | 0.149 |
| TOFusC l (Temporal Occipital Fusiform Cortex Left) | 0.185 | 0.552 | 0.117 |
| OFusG r (Occipital Fusiform Gyrus Right) | 0.204 | 0.552 | 0.138 |
| OFusG l (Occipital Fusiform Gyrus Left) | 0.197 | 0.585 | 0.118 |
| FO r (Frontal Operculum Cortex Right) | -0.178 | 0.186 | 0.148 |
| FO l (Frontal Operculum Cortex Left) | -0.087 | 0.355 | 0.177 |
| CO r (Central Opercular Cortex Right) | -0.105 | 0.421 | 0.135 |
| CO l (Central Opercular Cortex Left) | -0.126 | 0.317 | 0.124 |
| PO r (Parietal Operculum Cortex Right) | -0.126 | 0.211 | 0.136 |
| PO l (Parietal Operculum Cortex Left) | -0.13 | 0.301 | 0.125 |
| PP r (Planum Polare Right) | 0.059 | 0.486 | 0.176 |
| PP l (Planum Polare Left) | 0.026 | 0.381 | 0.158 |
| HG r (Heschl's Gyrus Right) | -0.037 | 0.3 | 0.127 |
| HG l (Heschl's Gyrus Left) | -0.057 | 0.308 | 0.125 |
| PT r (Planum Temporale Right) | -0.104 | 0.31 | 0.147 |
| PT l (Planum Temporale Left) | -0.065 | 0.429 | 0.131 |
| SCC r (Supracalcarine Cortex Right) | -0.028 | 0.237 | 0.116 |
| SCC l (Supracalcarine Cortex Left) | -0.013 | 0.24 | 0.112 |
| OP r (Occipital Pole Right) | 0.133 | 0.513 | 0.15 |
| OP l (Occipital Pole Left) | 0.121 | 0.449 | 0.163 |
| Thalamus r | 0.076 | 0.56 | 0.181 |
| Thalamus l | 0.067 | 0.573 | 0.157 |
| Caudate r | 0.012 | 0.495 | 0.153 |
| Caudate l | 0.058 | 0.411 | 0.149 |
| Putamen r | 0.06 | 0.392 | 0.162 |
| Putamen l | 0.093 | 0.534 | 0.156 |
| Pallidum r | -0.031 | 0.412 | 0.177 |
| Pallidum l | 0.144 | 0.538 | 0.154 |
| Hippocampus r | 0.151 | 0.504 | 0.168 |
| Hippocampus l | 0.1 | 0.545 | 0.141 |
| Amygdala r | 0.055 | 0.482 | 0.166 |
| Amygdala l | -0.007 | 0.458 | 0.175 |
| Accumbens r | 0.07 | 0.346 | 0.151 |
| Accumbens l | 0.064 | 0.387 | 0.153 |
| Brain-Stem | 0.044 | 0.571 | 0.16 |
| Cereb1 l (Cerebelum Crus1 Left) | 0.153 | 0.567 | 0.155 |
| Cereb1 r (Cerebelum Crus1 Right) | 0.228 | 0.684 | 0.17 |
| Cereb2 l (Cerebelum Crus2 Left) | 0.227 | 0.551 | 0.17 |
| Cereb2 r (Cerebelum Crus2 Right) | 0.291 | 0.632 | 0.164 |
| Cereb3 l (Cerebelum 3 Left) | -0.094 | 0.21 | 0.129 |
| Cereb3 r (Cerebelum 3 Right) | -0.023 | 0.326 | 0.169 |

| Region (Harvard Oxford Atlas, extended) | Mean *r* | Max *r* | Std *r* |
|---|---|---|---|
| Cereb45 l (Cerebelum 4 5 Left) | -0.02 | 0.36 | 0.137 |
| Cereb45 r (Cerebelum 4 5 Right) | -0.008 | 0.352 | 0.151 |
| Cereb6 l (Cerebelum 6 Left) | 0.079 | 0.507 | 0.169 |
| Cereb6 r (Cerebelum 6 Right) | 0.079 | 0.438 | 0.149 |
| Cereb7 l (Cerebelum 7b Left) | 0.133 | 0.579 | 0.236 |
| Cereb7 r (Cerebelum 7b Right) | 0.199 | 0.527 | 0.173 |
| Cereb8 l (Cerebelum 8 Left) | 0.053 | 0.49 | 0.158 |
| Cereb8 r (Cerebelum 8 Right) | 0.101 | 0.569 | 0.156 |
| Cereb9 l (Cerebelum 9 Left) | 0.073 | 0.478 | 0.175 |
| Cereb9 r (Cerebelum 9 Right) | 0.042 | 0.457 | 0.183 |
| Cereb10 l (Cerebelum 10 Left) | 0.165 | 0.483 | 0.16 |
| Cereb10 r (Cerebelum 10 Right) | -0.049 | 0.459 | 0.186 |
| Ver12 (Vermis 1 2) | -0.117 | 0.235 | 0.144 |
| Ver3 (Vermis 3) | -0.027 | 0.387 | 0.157 |
| Ver45 (Vermis 4 5) | -0.053 | 0.335 | 0.162 |
| Ver6 (Vermis 6) | 0.016 | 0.38 | 0.135 |
| Ver7 (Vermis 7) | 0.099 | 0.45 | 0.189 |
| Ver8 (Vermis 8) | 0.047 | 0.385 | 0.113 |
| Ver9 (Vermis 9) | 0.139 | 0.435 | 0.121 |
| Ver10 (Vermis 10) | 0.059 | 0.421 | 0.216 |
| Extended amygdala r | 0.093 | 0.218 | 0.075 |
| Extended amygdala l | -0.015 | 0.23 | 0.155 |
| Substantia nigra pars compacta r | 0.042 | 0.24 | 0.114 |
| Substantia nigra pars compacta l | -0.003 | 0.314 | 0.125 |
| Nucleus ruber r | -0.02 | 0.423 | 0.186 |
| Nucleus ruber l | 0.142 | 0.48 | 0.134 |
| Substantia nigra pars reticularis r | 0.041 | 0.264 | 0.1 |
| Substantia nigra pars reticularis l | 0.112 | 0.416 | 0.183 |
| Parabrachial pigmented nucleus r | -0 | 0.244 | 0.104 |
| Parabrachial pigmented nucleus l | 0.015 | 0.174 | 0.107 |
| Ventral tegmental area r | 0.095 | 0.223 | 0.143 |
| Ventral tegmental area l | 0.075 | 0.211 | 0.102 |
| Ventral pallidum r | -0.127 | 0.035 | 0.082 |
| Ventral pallidum l | -0.03 | 0.119 | 0.122 |
| Hypothalamus r | 0.029 | 0.382 | 0.145 |
| Hypothalamus l | 0.04 | 0.387 | 0.167 |
| Mammilary body r | -0.163 | -0.035 | 0.107 |
| Mammilary body l | -0.074 | 0.255 | 0.211 |
| Subthalamic nucleus r | 0.126 | 0.388 | 0.103 |
| Subthalamic nucleus l | 0.041 | 0.251 | 0.136 |
| NTS new r | 0.095 | 0.461 | 0.145 |
| NTS new l | -0.137 | 0.197 | 0.126 |
